# Dynamic responses to rejection in the transplanted human heart revealed through spatial transcriptomics

**DOI:** 10.1101/2025.02.28.640852

**Authors:** Kaushik Amancherla, Angela M. Taravella Oill, Xavier Bledsoe, Arianna L. Williams, Nelson Chow, Shilin Zhao, Quanhu Sheng, David W. Bearl, Robert D. Hoffman, Jonathan N. Menachem, Hasan K. Siddiqi, Douglas M. Brinkley, Evan D. Mee, Niran Hadad, Vineet Agrawal, Jeffrey Schmeckpepper, Aniket S. Rali, Stacy Tsai, Eric H. Farber-Eger, Quinn S. Wells, Jane E. Freedman, Nathan R. Tucker, Kelly H. Schlendorf, Eric R. Gamazon, Ravi V. Shah, Nicholas Banovich

## Abstract

Allograft rejection following solid-organ transplantation is a major cause of graft dysfunction and mortality. Current approaches to diagnosis rely on histology, which exhibits wide diagnostic variability and lacks access to molecular phenotypes that may stratify therapeutic response. Here, we leverage image-based spatial transcriptomics at sub-cellular resolution in longitudinal human cardiac biopsies to characterize transcriptional heterogeneity in 62 adult and pediatric heart transplant (HT) recipients during and following histologically-diagnosed rejection. Across 28 cell types, we identified significant differences in abundance in *CD4*+ and *CD8*+ T cells, fibroblasts, and endothelial cells across different biological classes of rejection (cellular, mixed, antibody-mediated). We observed a broad overlap in cellular transcriptional states across histologic rejection severity and biological class and significant heterogeneity within rejection severity grades that would qualify for immunomodulatory treatment. Individuals who had resolved rejection after therapy had a distinct transcriptomic profile relative to those with persistent rejection, including 216 genes across 6 cell types along pathways of inflammation, IL6-JAK-STAT3 signaling, IFNα/IFNγ response, and TNFα signaling. Spatial transcriptomics also identified genes linked to long-term prognostic outcomes post-HT. These results underscore importance of subtyping immunologic states during rejection to stratify immune-cardiac interactions following HT that are therapeutically relevant to short- and long-term rejection-related outcomes.

## INTRODUCTION

Solid-organ transplantation is the definitive treatment for end-stage organ failure. However, allograft rejection remains common early after transplantation (up to ≈40% of recipients experience rejection within 1-year post-transplant) and results in long-term graft failure and death^1–6^. Diagnosis of acute rejection and its response to therapy relies upon tissue histology^7^, subject to inter-observer variability^8^ that impacts precision of immunomodulatory therapies and potential for over- or under-immunosuppression^9–11^ with downstream clinical consequence. Heart transplantation (HT) represents a paradigmatic setting for this process: acute rejection affects ≈20% of recipients within the first-year post-HT^6^ and places patients at risk for chronic rejection phenotypes (cardiac allograft vasculopathy [CAV]) that limit long-term survival^12–14^. Rejection severity is graded by invasive endomyocardial biopsy (EMB)^9,15,16^, though some evidence suggests histology alone does not precisely phenotype rejection: (1) clinical presentation varies dramatically within the same grade of histologic rejection^17^; (2) response to anti-rejection therapy is heterogenous, including lack of response in some patients^18,19^. Moreover, there remains a substantial gap connecting acute rejection and its resolution to long-term outcomes (e.g., CAV). Given the routine performance of EMB for histologic rejection surveillance post-HT, HT represents a unique setting to study the impact of deeper spatially resolved molecular profiling that may inform precision approaches.

Here, we use spatial transcriptomics at sub-cellular resolution to delineate the molecular underpinnings of acute rejection in its different biological forms (cellular rejection/ACR, antibody-mediated rejection/AMR, mixed rejection) in hearts from 62 children and adults following HT. We studied spatial transcriptional profiles across rejection class and type as well as serial biopsies within the same patients during and after rejection therapy to map histologic-transcriptional concordance and dynamic response post-rejection treatment. Finally, we explored the relation of cell-specific gene expression signatures to CAV development to assess the ability of spatially resolved genomics to highlight genetic programs during and immediately following acute rejection relevant to long-term outcomes. Our overall goal was to create a large longitudinal biopsy resource in HT as a clinically relevant paradigm to study relevance of tissue-based, spatially-resolved genomics as an actionable molecular phenotype in acute rejection post-solid organ transplantation.

## RESULTS

### Study participants

Our study overview is outlined in **Fig. 1A**. We obtained longitudinal EMBs from 49 adult and 13 pediatric HT recipients. The median recipient age at the time of HT was 52 years (27% female) for adults and 6 years (62% female) for children. Complete donor and recipient characteristics are shown in **Table 1**. Biopsies were collected at a median of 313 days and 407 days post-HT for EMBs during rejection and following immunomodulatory therapies, respectively (**Supplementary** Fig. 1). Among biopsies demonstrating rejection, 67.7% comprised of acute cellular rejection (ACR), 11.3% antibody-mediated rejection (AMR), and 21% mixed cellular-antibody mediated rejection. Consistent with clinical practice, we observed a broad range of immunomodulatory therapies (corticosteroids, 50%; cytolytic T-cell targeted therapies, 11.3%; B-cell targeted therapies, 21%; combination cytolytic/B-cell therapy, 16.1%; no therapy, 1.6%; **Fig. 1B**).

**Figure 1.**
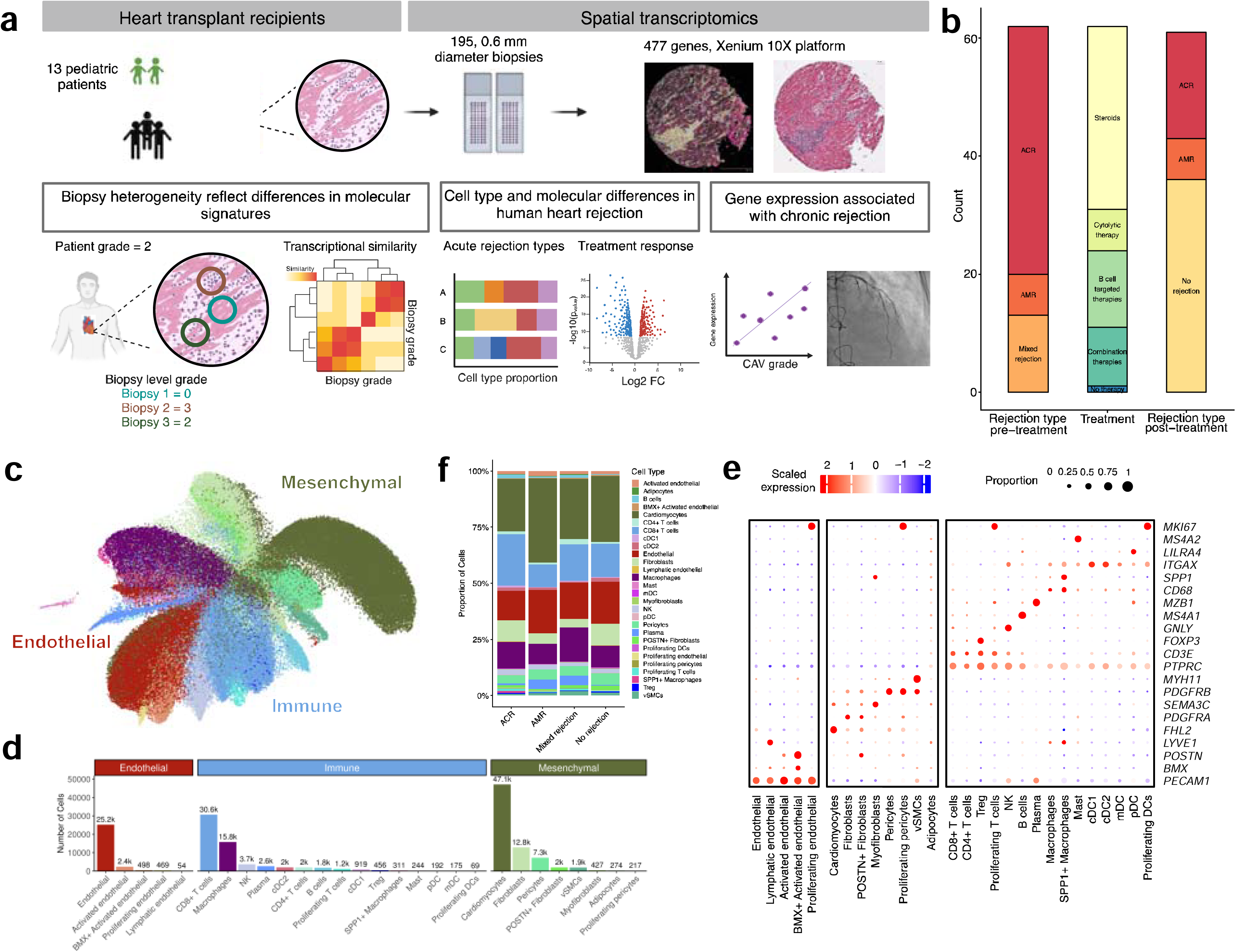
**(A)** Study overview. **(B)** Alluvial plot demonstrating rejection grading at the time of diagnosis in the left column, immunomodulatory therapies in the middle column, and biopsy grading following therapies in the right column. B-cell targeted therapies include combinations of intravenous immunoglobulin, plasmapheresis, rituximab, bortezomib, and eculizumab. Cytolytic therapies include thymoglobulin. Combination therapies correspond to combinations of cytolytic and B-cell targeted therapies. **(C)** Uniform manifold approximation and projection (UMAP) representing cell types identified. A total of 10,343,560 high quality transcripts were measured among 162,638 cells. **(D)** Bar plot demonstrating the 28 cell types identified and their respective cell numbers measured. **(E)** Cell type composition across rejection types. **(F)** Dotplot heatmap of select marker genes used for annotated cell types in the dataset. See extended data Fig. 1 for the expanded version.

**Table 1.** Baseline characteristics of the cohort.

| Characteristic | Overall, N = 62<sup>1</sup> | Age Group |  |
| --- | --- | --- | --- |
|  |  | Adult, N = 49<sup>1</sup> | Pediatric, N = 13<sup>1</sup> |
| Age at transplant (years) | 47 (25, 58) | 52 (44, 62) | 6 (2, 12) |
| Recipient gender |  |  |  |
| Female | 21 (34%) | 13 (27%) | 8 (62%) |
| Male | 41 (66%) | 36 (73%) | 5 (38%) |
| Class 1 DSAs | 22 (35%) | 14 (29%) | 8 (62%) |
| Class 2 DSAs | 40 (65%) | 27 (55%) | 13 (100%) |
| Deceased | 20 (32%) | 16 (33%) | 4 (31%) |
| CAV grade |  |  |  |
| 0 | 38 (61%) | 31 (63%) | 7 (54%) |
| 1 | 17 (27%) | 14 (29%) | 3 (23%) |
| 2 | 3 (4.8%) | 1 (2.0%) | 2 (15%) |
| 3 | 4 (6.5%) | 3 (6.1%) | 1 (7.7%) |
| Donor death type |  |  |  |
| DBD | 58 (94%) | 45 (92%) | 13 (100%) |
| DCD | 4 (6.5%) | 4 (8.2%) | 0 (0%) |
| Donor HCV NAT status |  |  |  |
| Negative | 50 (81%) | 37 (76%) | 13 (100%) |
| Positive | 12 (19%) | 12 (24%) | 0 (0%) |
| Donor age (years)<sup>2</sup> | 27 (20, 36) | 30 (23, 37) | 7 (2, 10) |
| Donor gender<sup>2</sup> |  |  |  |
| Female | 17 (28%) | 13 (27%) | 4 (33%) |
| Male | 43 (72%) | 35 (73%) | 8 (67%) |

<sup>1</sup>Median (IQR); n (%)
<sup>2</sup>Missing data on two individuals due to transfer of care from another transplant center.
DSA = donor-specific antibody; DBD = donation after brain death; DCD = donation after circulatory death; HCV = Hepatitis C virus; NAT = nucleic acid test.

### Compositional and spatial landscape of acute rejection highlights functionally relevant cell types post-HT

To better understand cell types driving allograft rejection, we evaluated compositional differences between the three types of rejection and following anti-rejection therapies. Across all samples, we measured 10,343,560 high quality transcripts among 162,638 cells comprising 28 cell types (**Fig. 1C**, **1D, and 1E**). Beyond canonical cardiac cell types (e.g., cardiomyocytes, fibroblasts, etc.), we identified *POSTN+* activated fibroblasts, myofibroblasts, *BMX+* activated endothelial cells, *SPP1+* macrophages, and proliferating pericytes, among others. Compared to prior single cell/single nuclei RNA-seq studies of the heart, we recover more immune and endothelial cells (**Supplementary** Fig. 2). Across annotated cell types, B cells, cardiomyocytes, *CD4+* and *CD8+* T cells, dendritic cells (cDC1 and cDC2), endothelial cells, fibroblasts, lymphatic endothelial cells, plasma cells, proliferating pericytes, proliferating T cells, and *SPP1+* macrophages were significantly differentially abundant in acute rejection (FDR < 0.1; **Fig. 1F and Supplementary Table 1**).

Overall, we observed cell types in rejection with plausible pathophysiologic connection related to its acute and chronic complications post-HT. In ACR, *CD8+* T cells, proliferating T cells, B cells, cDC1, fibroblasts, and *SPP1+* macrophages comprised the highest cell type proportion. Of note, fibroblasts and *SPP1*+ macrophages may mediate cardiac fibrosis^20,21^, a precursor finding of later CAV and allograft failure^22,23^. Moreover, *SPP1*+ macrophages have recently been implicated as key drivers of atrial fibrillation^24^, associated with acute rejection^25,26^. In contrast, we observed the highest proportions of endothelial cells, plasma cells, cardiomyocytes, macrophages, and proliferating pericytes in individuals with AMR, with a lesser proportion of active adaptive inflammatory cells (*CD4+*/*CD8+* T cells), with a trend toward increased activated and proliferating endothelial cells. Given the link between AMR, microvascular injury, and long-term CAV and graft failure, the high abundance of antibody-producing plasma cells, endothelial cells, and proliferating pericytes—all relevant to endothelial and coronary vascular injury—support the clinical epidemiology.

We next sought to establish the spatial localization of cell types in AMR and ACR. To this end, we calculated the pairwise likelihood of cell types being in close proximity (**Fig. 2**). In both AMR and ACR we find all subsets of T cells tend to be in close proximity to one another. In ACR we find this immune localization also includes pDCs and mDCs (**Fig. 2A**). Furthermore, within ACR biopsies, NK cells closely localize with endothelial, activated endothelial cells, and pericytes. In AMR, we find an additional localization of macrophage and fibroblast subtypes, where the fibroblasts are also in close proximity to BMX+ activated endothelial cells and proliferating pericytes (**Fig. 2B**). These results are consistent with observed focal fibrosis near damaged vasculature ^9–12,27^.

**Figure 2.**
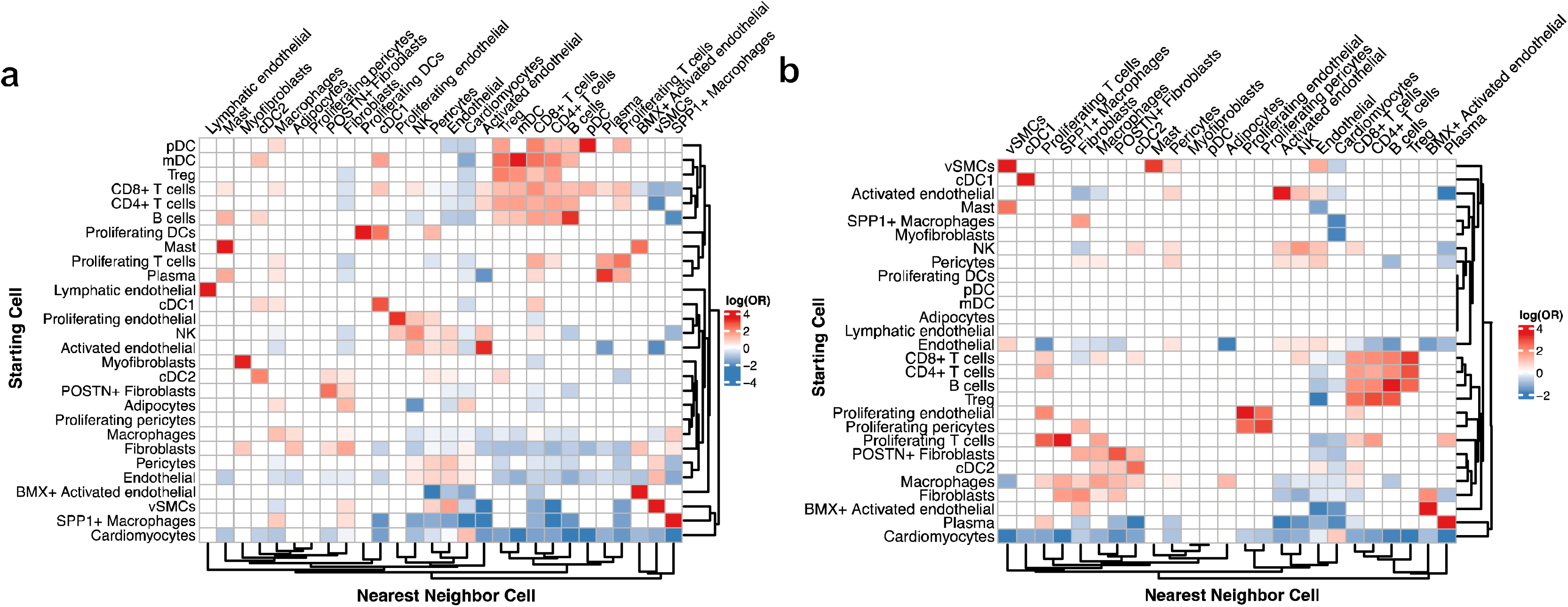
Cell proximity results across **(A)** ACR pre-treatment samples and **(B)** AMR pre-treatment samples. The heatmaps show the log odds ratio (log(OR)) where positive values (red) represent cell-type pairs that are proximal to each other while a negative value (blue) represent cell-type pairs that are less likely to be found near each other. Non-significant results (FDR > 0.05) are coded as 0.

### Significant transcriptional heterogeneity exists within and across histologic rejection grade

Histologic rejection grades drive clinical therapies post-HT^7,9–11^. However, despite similar appearances on histology, clinical presentation is heterogenous ranging from asymptomatic incidental findings on routine surveillance biopsy to cardiogenic shock. Similarly, therapeutic response is varied without predictability of histological response to immunomodulatory therapies. Therefore, we next sought to understand concordance between these clinically utilized histologic grades and transcriptional phenotypes within rejection. Clinical grading of EMBs to determine acute rejection grade was carried out according to International Society for Heart and Lung Transplantation (ISHLT) society standards^7,9^ by an experienced cardiac transplant pathologist (R.D.H) using whole-slide biopsies—clinical “gold standard”— distinct from those used for spatial profiling (tissue microarrays [TMA] constructed from biopsies). Notably, clinical grading requires multiple biopsies from the same patient as local pathologic remodeling is heterogenous. While this heterogeneity adds complexity to clinical rejection diagnosis, it also provides an opportunity to understand the degree to which local remodeling drives local cellular states. To this end, we generated high quality H&E images from samples after spatial transcriptomic analysis, enabling matched histology and molecular analysis. Using matched criteria to clinical grading, we generated a biopsy level grade for each pre-treatment ACR TMA sample (**Fig. 3A**). Next, we examined the transcriptional similarity of cells within and between biopsy grades. In general, we found cells from ACR grade ≤2R biopsies were more similar to other lower grade biopsies, with grade 3R biopsies having a distinct molecular profile (**Fig. 3B**). This association was particularly pronounced for fibroblasts, cardiomyocytes, and endothelial cells which were significant transcriptional similarity across nearly all low-grade biopsies and a distinct profile in grade 3R biopsies. Overall, these data suggest that the molecular state of cells are heavily influenced by the local remodeling regardless of the overall patient grade.

**Figure 3.**
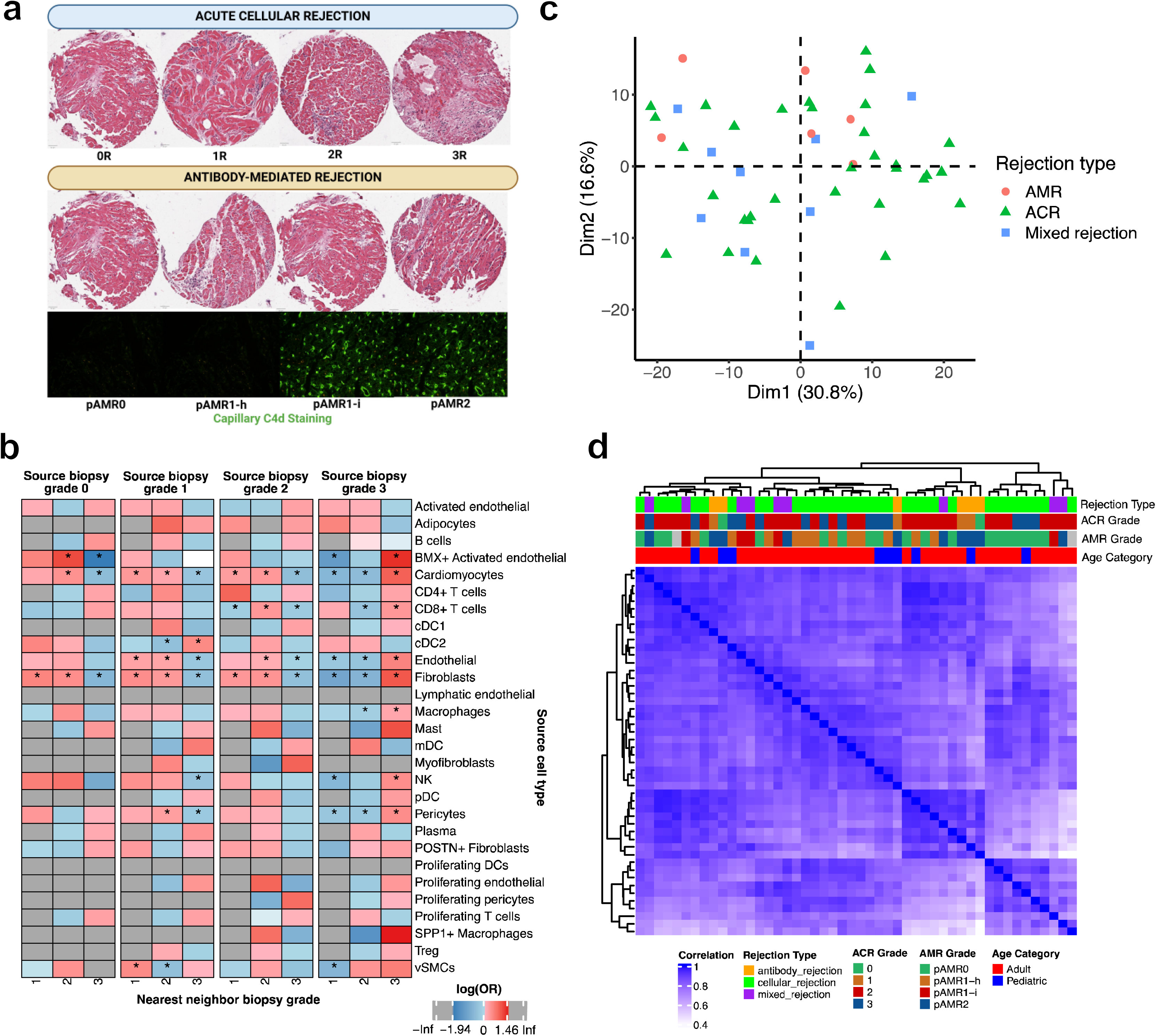
**(A)** Representative examples of rejection types and severity. Clinically-meaningful rejection grades warranting augmented immunomodulatory therapies include 2R and 3R within ACR and pAMR1-i and pAMR2 within AMR. This panel was made, in part, using Biorender. **(B)** Heatmap showing transcriptional similarity within and between biopsy grades. A positive log(OR) (red) represents biopsy grade pairs that are more transcriptionally similar, while negative log(OR) (blue) represents biopsy grade pairs that are less transcriptionally similar. Asterisks denote significant (FDR < 0.05) biopsy grade pair associations. **(C)** Principal components analysis (PCA) plot of all three rejection types based on pseudobulk gene expression demonstrates lack of clear clustering by histological rejection diagnosis. **(D)** Cluster dendrogram assessing similarity across rejection types. These results support that rejection is molecularly heterogeneous across biological class (e.g., ACR vs AMR) and within the same severity of rejection grading (e.g., grade 2R cellular rejection), despite similar histological appearance.

Turning our attention to the clinical (i.e. patient level) grading, rather than at the TMA level, we sought to quantify transcriptional heterogeneity across individuals. To this end, we created a pseudobulk expression profile for each individual and calculated the principal components (PCs) of gene expression to reduce dimensionality (**Supplementary** Figs. 3 and 4). The PC projections using histologically-determined grading according to clinical standards did not clearly separate the three types of rejection (**Fig. 3C; Supplementary** Fig. 5), suggesting underlying heterogeneity not explained by histology alone. This was reinforced by hierarchical clustering, which demonstrated broad overlap in gene expression profile across rejection types, grades, and patient ages (**Fig. 3D**). Together, these results suggest that molecular heterogeneity (masked by histology alone) may potentially explain the substantial variability in clinical presentation and therapeutic response that exists even within a given rejection grade.

### Spatial transcriptomics resolves functionally relevant patterns of gene expression between ACR and AMR

Clinical treatments for ACR and AMR largely differ, with ongoing uncertainty around selection of optimal therapies across rejection^7,9^. We hypothesized that cell-specific differences in gene expression between ACR and AMR (**Supplementary Table 2**) may shed light on clinical-hemodynamic differences between these types of rejection^9^. Between ACR and AMR, we identified 247 significantly differentially expressed (DE) genes across 15 cell types (FDR < 0.05; **Fig. 4A**). The cell type with the largest number of DE genes was *POSTN*+ fibroblasts (128 DE genes), followed by B cells (44 DE genes), and *CD8*+ T cells (32 DE genes; **Fig. 4B**).

**Figure 4.**
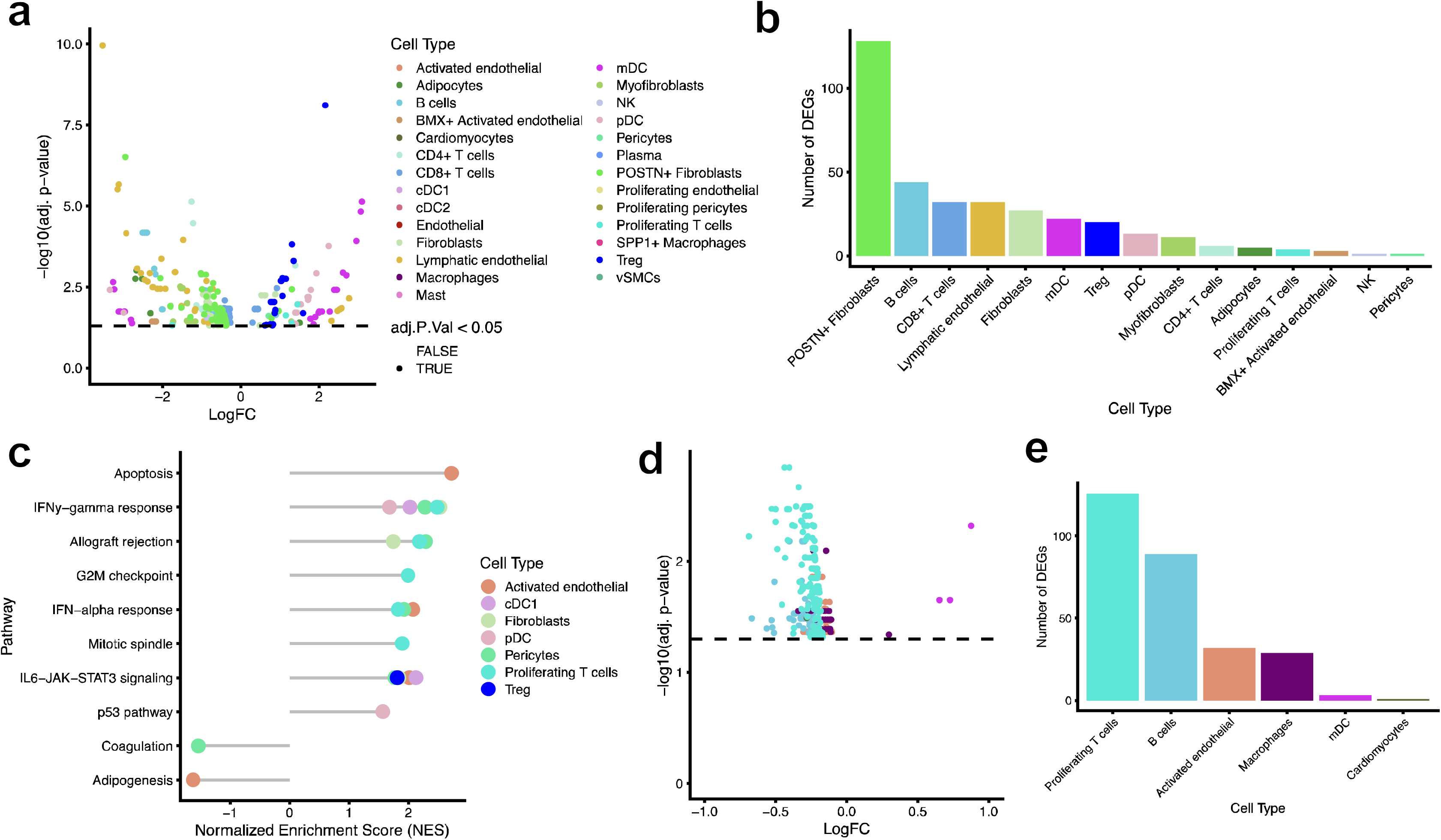
**(A)** Volcano plot representing differentially expressed (DE) genes between ACR and AMR. A positive log_2_FC means upregulated in ACR. A negative log_2_FC means upregulated in AMR. The color of each dot represents its corresponding cell type. Transparency reflects adjusted P-values, with transparent dots below the dashed black line representing those with adjusted P-values ≥ 0.05. **Supplementary Table 2** contains the full DE results. **(B)** Bar plot demonstrating the number of differentially expressed genes (DEGs) among cell types. **(C)** Gene set enrichment analysis (GSEA) depicting results of hallmark pathways that showed significance at an FDR < 0.1. A positive normalized enrichment score (NES) means enriched in ACR while a negative NES means enriched in AMR. **Supplementary Table 3** contains full results of the GSEA. **(D)** Volcano plot representing DE genes between responders and nonresponders to immunomodulatory therapies during ACR. A positive log_2_FC means upregulated in persistent ACR (nonresponders). A negative log_2_FC means upregulated in those whose ACR resolved (responders). The color of each dot represents its corresponding cell type. Transparency reflects adjusted P-values, with transparent dots below the dashed black line representing those with adjusted P-values ≥ 0.05. **Supplementary Table 6** contains the full DE results. **(E)** Bar plot demonstrating the number of differentially expressed genes (DEGs) among cell types.

Given the role *POSTN*+ fibroblasts have been show to play in mediating cardiac fibrosis^20^, these cells showed high expression of genes associated with fibroblast activation (e.g., *POSTN*^20^, *TLR4*^28^), myocardial ischemia and reperfusion injury (*S100A1*^29,30^, *SELE*^31^, *IL1B*^32^, *CCR7*^33^), antigen presentation (*HLA-DQB2*^34,35^,), and organ fibrosis (*TLR4*^28,36^, *CD5L*^37,38^, *CCR2*^39–41^, *CX3CR1*^42,43^, *CXCR4*^44^, *CXCL8*^45^) in patients with AMR. These pathways of vascular dysfunction and ischemia-reperfusion injury not captured by histology link transcriptional profiles within the rejecting cardiac allograft with long-term CAV. Proinflammatory genes (e.g., *CX3CL1*, *CXCL8*, *ICAM1*, etc.) were highly expressed in graft-infiltrating *CD4*+ and *CD8*+ T cells during both ACR and AMR non-specifically. Consistent with a dynamic role in acute inflammatory responses in immunologic injury, within Tregs we observed several DE genes that may limit inflammation (increased expression of *CXCL11*^46^, *FGL2*^47^, *IL2RA*^48^) and those that were linked to autoimmune disease (*CSF2RB*^49^). In particular, increasing expression of *IL2RA* functions may serve as a “sink” for locally-produced IL-2 to curb inflammation^48^. Of note, we also observed increased expression of *MYD88* in pDCs and *CD8*+ T cells during ACR, consistent with preclinical data suggesting that *MYD88* expression promotes acute rejection through recruitment of antigen-presenting cells and alloreactive T cells^50–53^.

In addition to inflammatory cell types, we also observed broad differences in cell types between ACR and AMR. We identified fibroblast-specific reduction in expression of *COL22A1* in ACR, linked to maintenance of vascular integrity^54^. *BMX*+ activated and lymphatic endothelial cells in AMR expressed genes associated with angiogenesis and endothelial function (*SNCG*^55^, *MMRN1*^56^, *PECAM1*^57^, *VWF*^58^, *PDGFRB*^59^). Gene set enrichment analysis (GSEA) using hallmark gene sets identified that upregulated genes in ACR are significantly enriched (FDR < 0.05) for IL6-JAK-STAT3 signaling in cDC1s and Tregs. Importantly, IFNγ signaling and genes involved in allograft rejection were enriched across multiple cell types (FDR < 0.05), including proliferating T cells and dendritic cells in ACR (**Fig. 4C**; **Supplementary Table 3**). These pathways of fibrosis, endothelial dysfunction, and loss of vascular integrity are implicated in long-term CAV development, linking transcriptional changes during ACR and AMR not clearly captured by histology to later mechanisms of graft failure and loss.

### Differentially expressed genes between ACR and AMR are not completely recapitulated in bulk studies of EMBs

To understand whether differential gene expression patterns identified in our study between ACR and AMR, a key diagnostic step that informs widely diverging immunomodulatory strategies, can also be identified in “bulk” studies (e.g., RNA-seq, microarray, etc.) we next sought to replicate our findings in a publicly-available EMB microarray dataset (GSE150059^60^). Comparing 76 patients with ACR and 179 patients with AMR, 58 shared genes (23.5%; out of 247 total unique DE genes identified in our study) were significantly DE between the conditions (**Extended Data** Fig. 2A and 2B). Of these, 20 genes (8.1%) shared the same positive directionality but only 12 genes (4.9%) shared the same negative directionality with our findings. There were 33 genes (13.4%) with discordant directionality. Importantly, 13 out of 33 discordant genes (39.4%) also shared overlap with the appropriate directionality within certain cell types. With <25% of spatially-resolved DE genes identified in the bulk data set and only ∼50% of them sharing the same directionality, these results suggest that bulk approaches may not fully recapitulate what single-cell and spatial studies capture and may be less informative for precision subphenotyping of allograft rejection. One confounding factor in bulk studies of EMBs is that, due to small biopsy size, histological confirmation of rejection within a research biopsy may not be feasible. As rejection is often patchy, this presents a limitation of solely using bulk approaches on EMBs.

### Spatial transcriptomics reveals differences in transcriptional states between resolution and persistence of rejection

Some patients experience ongoing rejection despite therapy, a unique opportunity to define molecular pathways of susceptibility to ongoing, “therapy resistant” rejection. In an effort to subphenotype this unique state, we examined cell-specific gene expression differences in those who had resolution of ACR following anti-rejection therapies (“Responders”) and those who did not (“Non-Responders”). We chose to prioritize ACR, given its higher frequency following HT and non-specificity of immunomodulatory anti-rejection therapy (e.g., corticosteroids, thymoglobulin, etc.)^6,10,11^. Of 32 individuals with ACR, 15 had resolution with anti-rejection therapies (17 did not). Among responders, we identified 16 unique differentially expressed genes across 7 cell types (FDR < 0.05; **Supplementary Table 4**). Despite resolution of rejection, post-treatment biopsies exhibited a significant increase in *SPP1* expression among *SPP1+* macrophages and *RAMP2* in pericytes, implicated in vascular integrity^61,62^. Among non-responders (individuals with ongoing rejection), we observed 14 DE genes across *SPP1*+ macrophages and dendritic cells despite anti-rejection therapy (FDR < 0.05; **Supplementary Table 5**). Importantly, there were no gene overlaps between those who responded to therapies compared with those who did not.

This lack of transcriptional overlap following immunomodulatory therapies between responders and non-responders led us to study pre-therapy biopsies (with rejection) between them to identify potential differences in the “baseline” gene expression profile that may mediate this difference. When comparing rejecting biopsies from responders vs. non-responders before anti-rejection therapy, we identified 216 unique differentially expressed genes (all FDR < 0.05) across 6 cell types (**Fig. 4D; Supplementary Table 6**). The overwhelming majority of DE genes were in proliferating T cells (126 genes), B cells (89 genes), activated endothelial cells (32 genes), and macrophages (29 genes; **Fig. 4E**). Surprisingly, except for 4 genes (*LYVE1* in macrophages; *DST*, *PECAM1*, and *SPIB* in mDCs), the remainder had significantly higher expression in ACR that was therapy responsive, suggesting lack of response in non-responders may owe to less inflammation at the time of initial rejection (despite a similar histologic grade).

Given the role of the endothelium in long-term CAV, we focused on activated endothelial cells in therapy-responsive ACR, identifying differential gene expression patterns consistent with inflammation (*AGER*^63,64^), immune system regulation (*HAVCR2*^65,66^), endothelial cell development (*LGR5*^67^, *SOX2*^68^), and maintenance of vascular integrity and angiogenesis (*SPP1*^69,70^, *ANPEP*^71–74^, *GPC1*^75^, *PPP1R1B*^76,77^, *HMGCS2*^78^, *FAS*^79^, *LEF1*^80^). Similarly, B cells—relevant to donor-specific antibody production important in alloreactivity—expressed genes associated inflammation regulation (*RETN*^81^, *GZMB*^82^, *FOSB*^83^, *IL1RL1*^84^, *CD40LG*^85,86^, *CD1E*^87^, *CX3CR1*^88^), proliferation and survival (*FOSB*^83^, *GDF15*^89^). Pathways involving genes related to allograft rejection, inflammation, IFNα/IFNγ and TNFα signaling, and IL6-JAK-STAT3 signaling were enriched predominantly vascular and supporting cells (e.g., endothelial cells, fibroblasts, pericytes) in responders (**Supplementary Table 7**). Taken together, these data support the hypothesis that substantial molecular differences exist in ACR despite similar histopathological findings and that individuals with higher baseline acute allograft inflammation may be more likely to respond to immunomodulatory therapies.

### Distinct cell-specific gene profiles are associated with incident cardiac allograft vasculopathy

Finally, to inform risk stratification for CAV based on molecular signatures and accelerate precision drug discovery to interrupt CAV pathogenesis following early immunological “hits” (often occurring years prior to overt CAV diagnosis), we related gene expression patterns during rejection and following therapy with CAV (**Fig. 5A and 5B**; **Supplementary Tables 8-12**). Across 14 cell types during ACR, we identified 116 unique genes associated with CAV. These included associations with CAV across pathways of angiogenesis (*ADGRL2*^90,91^ in VSMCs), vascular inflammation (*CX3CL1*^92^ and *ITGAM*^93^ in VSMCs; the BET genes [e.g., *BRD2*, *FGL2*^94^, *MEF2C*^95^, the multimerins^56,96^, *PLA1A*^97^, *VCAM1*^98^, and *CFHR3*^99^ in *BMX+* activated endothelial cells; the TNF receptor superfamily members^100^ in endothelial cells) and proliferation (*FAS*^101–103^ in VSMCs), and pro-coagulation (*CD40*^104^ in *BMX+* activated endothelial cells). These pathways—activated during ACR—have been implicated in CAV. Several genes have been previously identified in microarray studies of biopsies during acute rejection, including *PLA1A*, *GGH*, *LYZ*, *CSF2RB*, *CX3CL1*, *NOMO1*, *MS4A3*, *UBD*, *MINAR2*, and *FCGR3A*^97,105^. Here, we show for the first time that these genes are expressed in a cell-specific fashion during acute rejection and are associated with CAV.

**Figure 5.**
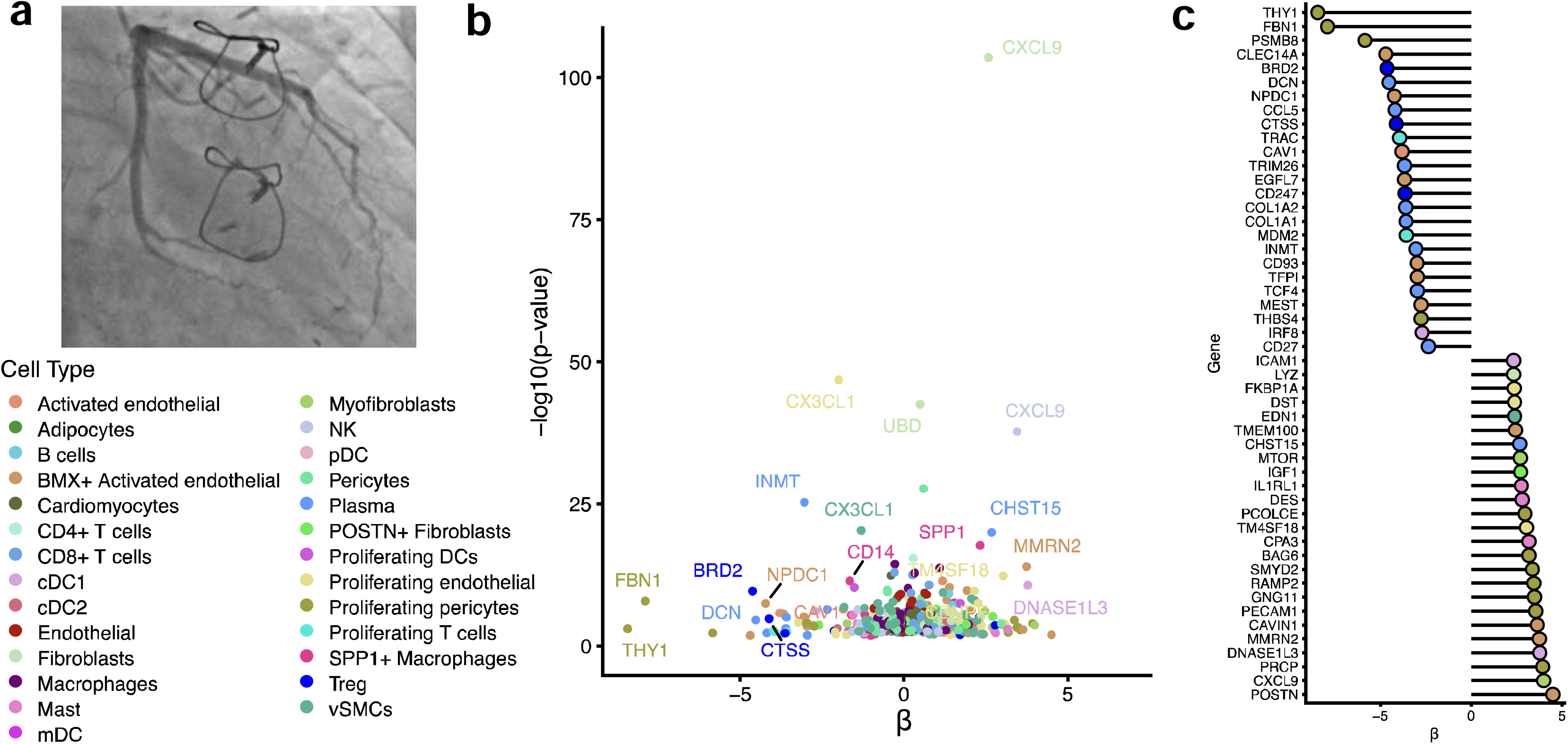
**(A)** Coronary angiogram of a patient included in this study with moderate CAV (CAV grade 2). **(B)** Association between cell-specific gene expression across biopsies spanning all three types of rejection and following immunomodulatory therapies. The x-axis represents the β from the generalized linear mixed model (GLMM) while the y-axis represents the -log_10_(P-value). The color of each dot represents its corresponding cell type. Transparency reflects adjusted P-values, with transparent dots representing those with adjusted P-values ≥ 0.05. **Supplementary Tables 8 through 12** contain the full results of the GLMM models across all biopsies. **(C)** The top 50 unique genes (arranged by absolute value of their β in the GLMM models) and cell types within those top betas are depicted.

Following anti-rejection therapies in ACR, which are expected to attenuate myocardial inflammation, we then tested for the presence of “residual” cell-specific inflammation that may be associated with CAV. We identified 83 unique significant genes, across 20 cell types, that were associated with CAV. None of these cell-specific genes were shared with those identified in ACR biopsies preceding therapies, suggesting the presence of evolving molecular processes despite histological resolution. The largest number of associations were in cell types implicated in CAV: those of the microvasculature (pericytes), supporting cells (fibroblasts), and antibody-producing cells (B cells and plasma cells). The strongest associations with CAV included *COL1A2* in proliferating endothelial cells and *MTOR* and *DES* expression in myofibroblasts. Currently, mTOR inhibitors are part of a limited set of therapies in the armamentarium against CAV. Our findings of early *MTOR* expression following rejection treatment specifically in myofibroblasts, previously shown to drive fibrosis and vasculitis^106–108^, may offer an opportunity for early interruption of CAV pathogenesis following rejection. Furthermore, GSEA identified enrichment of pathways involved in endothelial-to-mesenchymal transition among cell types of the vasculature (pericytes, VSMCs, fibroblasts; FDR < 0.05; **Supplementary Table 13**), which we have previously identified in the myocardium of individuals with severe CAV^27^. These results show that despite anti-rejection therapies, residual inflammatory risk (unmeasured by histology alone) persists and is associated with implications for long-term graft health.

Among patients with AMR, 73 genes across 17 cell types were associated with CAV. Significant gene associations predominantly spanned microvasculature and supporting cell types (VSMCs, pericytes, activated endothelial cells, and fibroblasts). These include genes associated with vascular integrity and angiogenesis (*FAS*^103,109^ in activated endothelial cells; *PECAM1*^57^, *SMYD2*^110^, *PCOLCE*^111^, and *RAMP2*^61^ in proliferating pericytes; *FBN1*^93,112^, *ACTG2*^113^, and *OGN*^114^ in VSMCs), fibroblast and complement activation (*BRD2*^115^, *C4A*^9,116^), and inflammation (*IRF8*^117^, *ICAM1*^118^, and *IL6R*^119^ in cDC1; *ITGAM*^120^ and *SLAMF7*^121^ in NK cells), among others. Following therapies, there were 155 genes broadly distributed across 17 cell types that remained associated with CAV. These included 37 shared genes but expressed across different cell types. The overwhelming majority of gene associations following AMR treatment were concentrated in cardiomyocytes, microvasculature, *CD8*+ T cells, and macrophages. These included genes related to endothelial integrity and inflammation (*PECAM1*^57^, *CD93*^122^, *FBN1*^112,123^, *TCF4*^124^, *VCAN*^21^, *MTOR*^106–108^ across subsets of endothelial cells), fibrosis and protection against heart failure (e.g., cardiomyocyte-specific increase in *CXCL10*^125–127^ expression to minimize fibrosis and angiogenesis, along with decrease in *TLR4*^128,129^ expression to protect against graft dysfunction). As expected, persistent inflammation across a variety of immune cells was prominently associated with CAV. Similar gene associations between mixed rejection and CAV were also identified, including 20 and 26 shared genes with AMR and ACR, respectively. The top 50 unique transcripts with the largest association (absolute value of β) and their cell type links are shown in **Fig. 5C**.

### Spatial transcriptomics identifies genes altered during acute rejection that are dysregulated during end-stage, severe CAV in transplanted human myocardium

To evaluate whether these genes were expressed in a similar cell-specific fashion in the myocardium following overt CAV development, we turned to our previously published single-nuclear RNA-sequencing data set that sampled left and right ventricular tissue from the human heart in CAV^27^. Across all rejection biopsies (including post-treatment ones) in our study, 309 unique genes were associated with CAV (FDR <0.05 for all). We prioritized these 309 genes and observed enrichment of gene expression in an expected cell-specific fashion (e.g., high expression of *PECAM1*, *VCAM1*, *ICAM1*, *CD93*, *POSTN*, etc., in endothelial and endocardial cells; *MYD88* in dendritic cells and monocytes; *IL6R*, *ITGAX*, *IRF8* across multiple immune cell types; among others; **Extended Data** Fig. 3, **Supplementary** Figure 6). These data suggest that cell-specific gene expression during rejection and following treatment—particularly genes involved in residual inflammation, angiogenesis/vascular integrity, and anti-fibrotic responses— are significantly expressed in a similar cell-specific fashion in end-stage CAV myocardium, transcriptionally linking “early” immunological phenomena to long-term CAV risk.

## DISCUSSION

Tissue-based molecular phenotyping is *sine qua non* of modern precision genetic approaches across a variety of disease states^130^. Penetrance of tissue genomic methods to subphenotype disease has been especially fruitful in diseases where cell-level heterogeneity is the rule (e.g., oncology^130–132^). While most chronic diseases do not lend themselves to tissue genetic-driven therapy, solid-organ transplantation represents a unique setting where histology may be routinely obtained during allograft rejection to drive therapy^7,9^. Unfortunately, variability in histologic interpretation and lack of individualization of therapies based on cellular transcriptomic states challenges precise application of immunomodulatory therapy (and downstream immunosuppression-related chronic clinical effect). Studies that directly examine tissue molecular transcriptomic states with reference to histology and dynamic changes with therapy across different modes of rejection are urgently needed to translate genomic advances enjoyed by oncology to this space.

Here, we answer this call by conducting spatial transcriptomics in 62 longitudinal adult and pediatric cardiac biopsies to characterize acute rejection following heart transplantation, dynamic responses to immunomodulatory therapies, and link cell-specific gene expression patterns to the development of chronic rejection (i.e., CAV). Our principal findings show (1) a diverse array of cell types are involved in acute rejection and response to immunomodulation, with *CD8+* T cells, proliferating T cells, B cells, cDC1, fibroblasts, and *SPP1+* macrophages comprising the highest proportion of cell types in ACR and endothelial cells, plasma cells, cardiomyocytes, and proliferating pericytes comprising the highest proportion of cell types in AMR. (2) There was substantial molecular heterogeneity both across rejection types and within the same rejection grades, masked by histology alone. Differences in gene expression patterns between AMR and ACR were only partially captured in an external microarray study of EMBs during rejection. (3) Baseline rejection biopsies are comprised of distinct transcriptomic profiles between patients who go on to respond to immunomodulatory therapies versus those who do not. (4) Cell-specific gene expression patterns both during rejection and following immunomodulatory therapies are associated with chronic rejection (i.e., CAV). These genes showed similar cell-specific expression patterns in an external snRNA-seq dataset of myocardium obtained from patients with end-stage CAV. Our findings link molecular heterogeneity with clinical heterogeneity and advance our knowledge of how acute rejection may mediate CAV. These are highly relevant clinical processes that are incompletely understood at this time. Furthermore, this unique dataset is available to the larger scientific community and may inform novel approaches to diagnosis of acute rejection, prediction of therapeutic response, and risk stratification for CAV development.

The current gold standard for rejection diagnosis remains histology supplemented with limited immunofluorescence or immunohistochemistry^7,9^. However, inter-pathologist variability in the grading of rejection is substantial^8^. More importantly, histology alone has failed to explain heterogeneity in clinical presentation, whether individuals will respond appropriately to anti-rejection therapies, and in risk stratifying for future CAV development. Using PCA and hierarchical clustering, we show that substantial molecular heterogeneity is present both across different types of rejection (ACR, AMR, mixed) and within the same grades of histologic rejection (e.g., during grade 2R ACR). This latter finding is particularly important clinically, as it may help explain why a spectrum of clinical presentations – spanning complete lack of symptoms to florid cardiogenic shock – exists within the same histologic grade of rejection and why some individuals have resolution of rejection with minimal intervention (e.g., outpatient augmentation of oral prednisone) while others require broadly-targeting T- or B-cell depleting therapies (e.g., thymoglobulin, intravenous immunoglobulin, plasmapheresis, rituximab, etc.).

Furthermore, while multiple observational studies have shown that acute rejection is a major risk factor for CAV development^13^, it remains incompletely understood as to how episodes of acute rejection are linked to CAV. While animal models have suggested that IFNγ signaling and interactions between both lymphoid and myeloid cells with coronary endothelial cells and VSMCs may play important roles ^133,134^, direct clinical translation has been limited by lack of clinical levels of immunosuppression. Our results in human cardiac tissue at the time of acute rejection highlight a number of cell-specific genes associated with CAV. Notably, the majority of significantly-associated genes are in cells of the vasculature (endothelial cells, *BMX+* activated endothelial cells, pericytes) and supporting cells (fibroblasts, VSMCs) and span across pathways angiogenesis (*ADGRL2*^90,91^ in VSMCs), vascular inflammation and proliferation (*CX3CL1*^92^ and *ITGAM*^93^ in VSMCs; the BET genes [e.g., *BRD2*], *FGL2*^94^, *MEF2C*^95^, the multimerins^56,96^, *PLA1A*^97^, *VCAM1*^98^, and *CFHR3*^99^ in *BMX+* activated endothelial cells; the TNF receptor superfamily members^100^ in endothelial cells; *FAS*^101–103^ in VSMCs), and promotion of coagulation/complement fixation. These microvasculature-directed transcriptomic changes suggest the potential for spatial molecular phenotyping approaches in risk stratifying individuals for future CAV. Importantly, CAV-associated genes that we identify in our study (“early” in disease course) are expressed in a similar cell-specific fashion in myocardium from patients with end-stage CAV requiring re-do HT.

As spatial technologies evolve to capture the whole transcriptome at subcellular resolution, prioritizing study of longitudinal EMBs may further elucidate novel insights into divergent cell-specific fates in those with healthy allografts in comparison to those who develop rejection or challenges to long-term graft survival (e.g., CAV). The few studies performed to date have largely prioritized “bulk” approaches (mostly microarray studies) to link molecular patterns with pathologist-given histologic grade^60,97,135–140^. These studies have been challenged by lack of cellular/subcellular resolution (“washing out” cell-specific gene expression patterns), inadequate resolution of the coronary microvasculature only accessible in modern spatial techniques (relevant to CAV), among other limitations. Data generated through approaches such as ours (e.g., spatial and single-cell transcriptomics) during a variety of therapies and long-term immunomodulatory strategies (e.g., mTOR inhibition) may substantially inform early biomarker and drug discovery to meaningfully prolong allograft survival in the modern era^141,142^.

Key strengths of this study include a large sample size spanning adult and pediatric HT, longitudinal human cardiac biopsies from the same patients, and modern spatial transcriptomics at sub-cellular resolution to interrogate cell-specific gene expression patterns. Furthermore, cardiac biopsy samples in our study were obtained from individuals without end-stage disease, which carries a very distinctive transcriptomic profile that may not reflect early, modifiable mechanisms^20,41,143–146^. Our study of rejection and its dynamic therapeutic response in non-end-stage allografts offers novel insights into immune-cardiac interactions that are not influenced by advanced disease. Nevertheless, our study does have important limitations. Despite sampling biases inherent in modern endomyocardial biopsy techniques, invasive biopsy remains the current gold-standard of clinical care post-HT. To limit variability, we used a single, experienced cardiac pathologist to review whole slide gradations of rejection, as well as TMA-specific gradation. Despite our sample size and longitudinal sampling, additional gene discovery in larger samples is warranted to expand mechanistic implication. In addition, our approach (Xenium) uses a targeted probe set informed by our prior experience and published literature. Nevertheless, our findings are biologically plausible with replication in single nuclear data. We anticipate that as technologies advance in conduct and economy, unbiased whole transcriptomics at sub-cellular resolution will substantially add to our findings.

In conclusion, our study establishes significant molecular heterogeneity across and within rejection unapparent by histology alone in adult and pediatric HT. Further, we identify distinct cell-specific gene expression patterns, only partially recapitulated in bulk studies, across dynamic responses to therapy using longitudinal cardiac biopsies. Finally, we link cell-specific gene expression to highly-relevant clinical outcomes (i.e., chronic rejection [CAV]) and show similar cell-specific gene expression in end-stage CAV. Together, our findings shed new insight into cardiac allograft rejection and provide a valuable resource for the larger transplant scientific community. We anticipate this work to be a step toward precision approaches to solid-organ transplantation.

## Methods

### Human samples and study approval

The overall study design is depicted in **Fig. 1A**. Formalin-fixed paraffin-embedded (FFPE) EMB samples were obtained from adult and pediatric HT recipients at Vanderbilt University Medical Center (VUMC) undergoing routine surveillance or for-cause endomyocardial biopsies between December 2017 and January 2023. Rejection grading was performed by clinical pathologists and confirmed by pathological review by an experienced cardiovascular transplant pathologist (R.D.H.) at both the whole slide and tissue microarray (TMA) levels to limit spatial variability. Only biopsies exhibiting clinically-relevant acute rejection were selected, along with post-treatment biopsies, and were defined according to accepted standards – ACR grade ≥2R was defined as ACR and pAMR1-i or pAMR2/3 were defined as AMR. The presence of both ACR and AMR concurrently on the same EMB was defined as mixed rejection. This study was approved by the VUMC institutional review board (IRB #200551).

### Tissue microarray construction

To minimize batch effect and improve run efficiency, multiple samples were placed on a single Xenium slide (available sample positioning area measuring 10.45 mm x 22.45 mm) using a tissue microarray (TMA). Slides of each FFPE EMB block were H&E stained for clinical diagnosis and were evaluated by a cardiovascular pathologist (R.D.H.), who selected two regions of interest per block, inclusive of the area of rejection. In EMBs without clinical rejection, two normal regions were selected. TMA blocks were constructed to target two 0.6mm cores per FFPE block and a total of 233 cores underwent sectioning at 5 µm thickness before being placed onto 2 Xenium slides. After drying overnight, the slides were placed in a sealed desiccator and stored at room temperature for ≤7 days.

### Xenium workflow

This workflow uses targeted gene panels that can be custom designed. A total of 477 unique genes were included in this study. Of these, 377 were in a pre-designed panel (Human Multi-Tissue and Cancer), which include markers of a large panel of immune cells as well as cells comprising the human myocardium. An additional 100 custom genes were selected, informed by published single-nuclear RNA-seq^27^, and bulk RNA-seq and microarray studies^60,97,137–140^. The full list of genes is available in **Supplementary Table 14**.

As part of sample preparation, all workstations and equipment were cleaned using RNAse Away (RPI 147002) and 70% isopropanol. Following placement on Xenium slides, tissue samples were deparaffinized and decrosslinked to access mRNA. Hybridization using the gene panel probes occurred for 18 hours at 50LC, followed by washing of unbound probes. Next, probes were ligated at 37LC for 2 hours followed by rolling circle amplification for 2 hours at 30LC. The slides were then washed, stained with the 10X Genomics commercial kit for improved cell segmentation, and background fluorescence (present in a number of tissues, including heart) was quenched chemically. Additional washes with PBS and PBS-T were undertaken, nuclei were stained using DAPI, and slides were loaded onto the Xenium Analyzer instrument.

Following loading of consumable reagents and the two Xenium slides, the Xenium Analyzer instrument decodes subcellular localization of mRNA targets in a fully automated fashion. After binding of fluorescently labeled oligos, samples underwent multiple cycles of hybridization, followed by acquisition of images and probe stripping are undertaken with images taken at 4240 x 2960 pixel field of views (FOVs). As each gene in the panel has a unique fluorescence pattern across image channels, focal areas of intensity were decoded and labeled according to the gene ID. All image FOVs and transcripts were stitched together using the DAPI-stained image as the anchor, followed by quality control (QC) for each detected transcript (instrument software version v1.1.2.4 and xenium-v1.1.0.2). Following the run on the Xenium Analyzer instrument, the chemical quencher was removed and slides underwent H&E staining. Histology images were taken using the 20X Leica Biosystems Aperio CS2 instrument. Cell segmentation was performed using the 10X Xenium software. All analyses were performed at the cell level.

The Xenium images and H&E images were integrated using ImageJ (v 2.14.0) based on identifiable landmarks on both images. Registered images were manually reviewed for any potential artifacts resulting from this process. In the Xenium Explorer, we used the lasso function to obtain cell IDs corresponding to each visible core/punch. Of the 233 targeted cores, 195 punches were lassoed and used in downstream analyses. The output files include coordinates in 2D space, corresponding gene, assigned cell, and quality metrics. Using this output, low quality transcripts (qv < 20) and transcripts that corresponded to blank probes were removed. Subsequently, Seurat (v5.0.1) was used and cell level count data was loaded into RStudio using Seurat’s LoadXenium function with default settings. Cells outside of lassoed regions were excluded and cells with 0 counts were removed from subsequent analyses, leaving a total of 189,109 cells. Cells were retained based on ≥ 12 transcripts corresponding to ≥ 10 unique genes, percentage of high-quality transcripts corresponding to blank probes ≤ 5, and nucleus area ≥ 6 and ≤ 70 µm. A total of 164,165 cells remained after filtering.

Dimensionality reduction and clustering were performed using a Docker container with RAPIDS (v21.8.1), which implements Scanpy (v1.8.1), while Seurat (v5.0.1) was used for downstream analyses and visualizations. RNA counts matrices underwent log1p normalization. Principal components analysis (PCA) was used for dimensionality reduction. Nearest neighbors distance matrix and a neighborhood graph was computed using 20 PCs and 20 nearest neighbors and was used to generate a uniform manifold approximation and projection (UMAP). Clustering was performed using the Leiden algorithm. Cell types were annotated using a similar approach as our prior work^147^, where first pass annotations were implemented to define the main lineages and a second pass was performed for fine level granular annotations. To identify cell types, we assessed expression of canonical marker genes and top markers using Presto (v1.0.0) wilcoxauc and top_markers functions (**Extended Data** Fig. 1). This resulted in 28 cell types including 5 endothelial, 15 immune, and 8 mesenchymal. Post annotation, we excluded cells from one patient that was not related to this study from subsequent downstream analyses, leaving a total of 162,638 cells.

### Comparison to scRNA human heart atlas

To compare lineage-level cell type recovery in our data, we compared our results with the human Heart Cell Atlas v2 single-cell/nuclei RNA and ATAC-seq data set^143^. We included all samples from the Heart Cell Atlas (HCA; 25 adult donors, 14 females and 11 males). Since the HCA data set comprised of unaffected donors and our data set was from heart transplant patients, we separated our biopsies into two groups – biopsies without evidence of rejection (normal myocardium) and biopsies with rejection – when comparing lineage proportions rather than aggregating all cells for each patient. For the HCA data, the proportion of cells from each lineage was calculated for each donor, while for our data, the proportion of cells from each lineage was calculated for each biopsy.

### PCA

To assess the heterogeneity across acute rejection we performed PCA using the factoextra R package. PCA was performed on matrices of cell type proportions and on pseudo-bulked expression for pre-treatment samples. Cell type proportions were calculated for each biopsy. For the expression PCAs, the log1p normalized data was averaged per gene per patient across all cell types and for each cell type separately.

### Transcriptional similarity analysis

To characterize the molecular profiles of endomyocardial biopsies of different rejection grades we stratified pre-treatment biopsies from ACR patients by grade and identified the closest cells in terms of expression. We ran RunPCA() and FindNeighbors() in Seurat for all cells from the same biopsy grade and for each cell type separately (source cell type). Then the nearest neighbor graph used to select the cell with the most transcriptional similarity to the source cell. A Fisher’s exact test was then performed for each source biopsy grade, by grouping source cells of the same biopsy grade and not, and if the closest transcriptional neighbor was from the same biopsy grade as the source cell or not. P-values were adjusted using FDR correction method.

### Cell type proximity analyses

To assess physical proximity between cell types we used the approach established in Vannan et al. 2025. Briefly, distance and angular direction were calculated between each cell and its neighboring cell within a 60-µm radius circle. The probability of cell types being proximal to one another was assessed using Fisher’s Exact by assigning cells to binarized proximal and non-proximal categories. P-values were adjusted using FDR correction method.

### Cell type proportion analyses

Cell type proportion differences were statistically assessed among AR types using the propeller method within the speckle R package^148^. Using the propeller default function, cell type proportions were calculated at the biopsy level for all pre-treatment biopsies, proportions were transformed using the “logit” method, and an ANOVA was performed.

### Differential gene expression (DGE) analyses

We performed cell-type specific DGE analyses using the limma voom framework^149,150^. For the DGE analysis comparing ACR and AMR pre-treatment and ACR patients that had resolution pre- vs post-treatment we pseudo-bulked on patient level. For DGE analysis pre-treatment only persistence vs resolution we pseudo-bulked on biopsy and regressed out biopsy grade since there is a difference in the distribution of biopsy score in pre-treatment resolved vs pre-treatment persistence (**Supplementary** Fig. 7). Pseudo-bulking was performed by averaging the log1p normalized data per gene and per patient or biopsy for each cell type separately. We removed cell types where there were less than 3 samples and kept genes that have expression in at least 3 samples. GSEA was performed using the fgsea and msigdbr packages, with a focus on Hallmark gene sets.

### Generalized Linear Mixed Models (GLMMs)

We performed a series of GLMMs to assess whether expression is associated with outcome (CAV grade) in ACR patients. GLMMs were performed using glmmTMB package in R. Patient age was added as a fixed effect and patient ID was added as a random effect. We pseudo-bulked per biopsy by averaging the log1p normalized data per gene and per biopsy for each cell type separately. Genes were removed from analysis if there were less than 5 samples with expression. P-values were adjusted using FDR correction method.

### Validation of DGE results with publicly-available microarray data set

GeoQuery was used to download microarray data (GSE150059)^60^. Data were first evaluated for quality and gene distribution. Subsequently, only biopsies labelled “ABMR” (representing AMR) and “TCMR” (representing ACR) were included for downstream analyses. The *limma* pipeline (lmFit) was used for DGE analyses. Multiple testing correction was performed by the Benjamini-Hochberg correction. Genes were considered significantly differentially expressed if BH-corrected P-value < 0.05 and absolute(log_2_FC) > 1.5.

### Validation of CAV-associated gene expression in single-nuclear RNA-sequencing of end-stage CAV myocardium

Prior data from our group focused on single-nuclear RNA-sequencing of cardiac biopsies from end-stage CAV patients undergoing re-do transplant and non-CAV controls was utilized, with data processed as previously described^27^. As CAV tissue was sampled from both the right (RV) and left ventricles (LV), we prioritized RV samples to maintain concordance with our current study (as EMBs are from the RV). Normalized gene expression outputs were used for visualization.

## AUTHOR CONTRIBUTIONS

A.M.T.O., K.A., X.B., S.Z., and N.B. performed analyses. K.A. collected all data and biopsies. R.D.H. evaluated biopsy rejection grading. K.A., R.V.S., and N.B. supervised the analyses. K.A., A.M.T.O., R.V.S., and N.B. wrote the manuscript. All authors interpreted data and edited the manuscript.

## DATA AVALIABILITY

Raw and processed data are deposited on GEO under accession GSE290577.

## CODE AVALIABILITY

Code relating to the analyses for this paper are available on GitHub at https://github.com/Banovich-Lab/Spatial_Heart_Rejection.

---

**Extended data Fig. 1.**
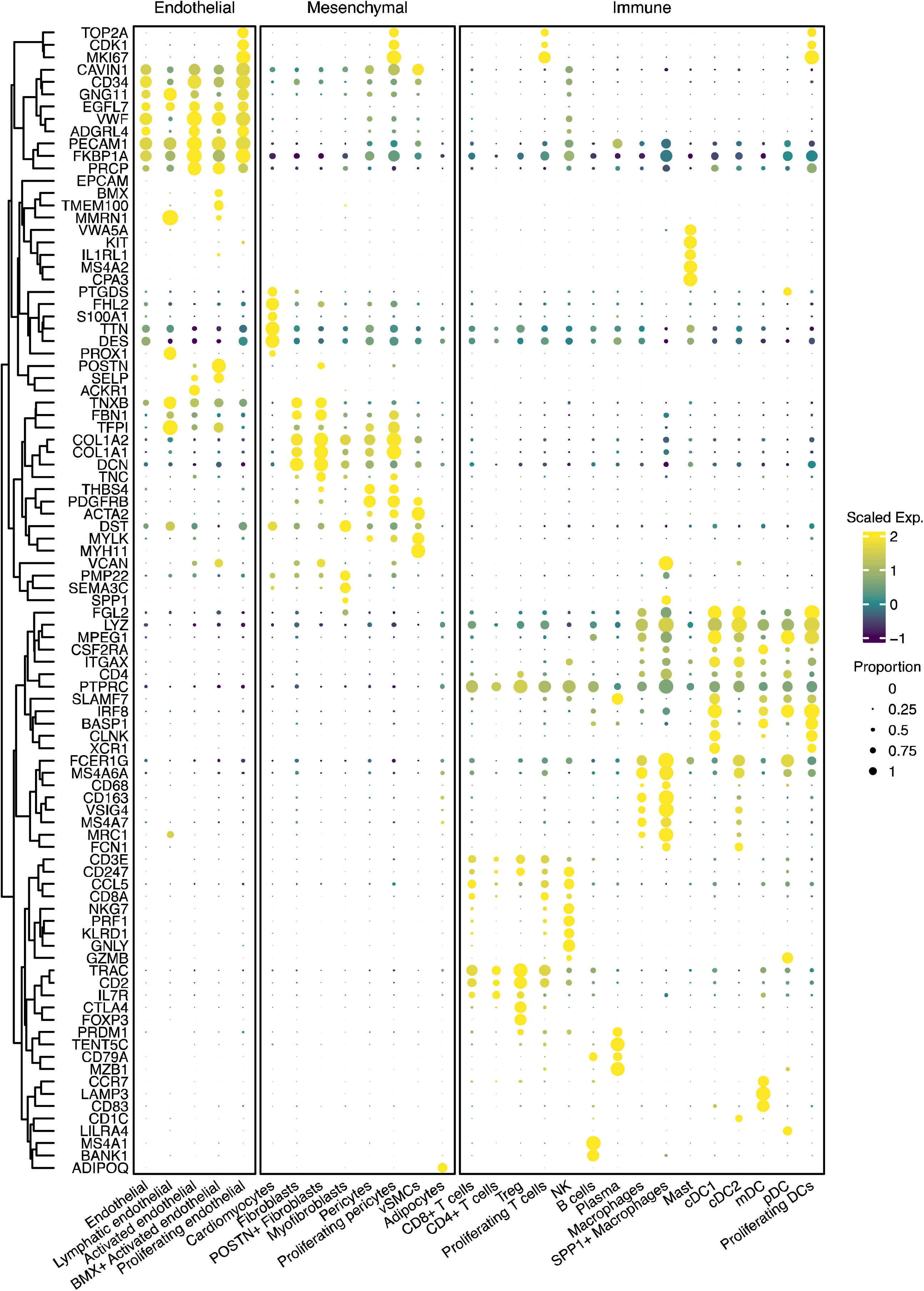
Marker genes for all cell types. A combination the top five expressed marker genes for each cell type and canonical marker genes are shown here to demonstrate cell type annotation in our data set.

**Extended data Fig. 2.**
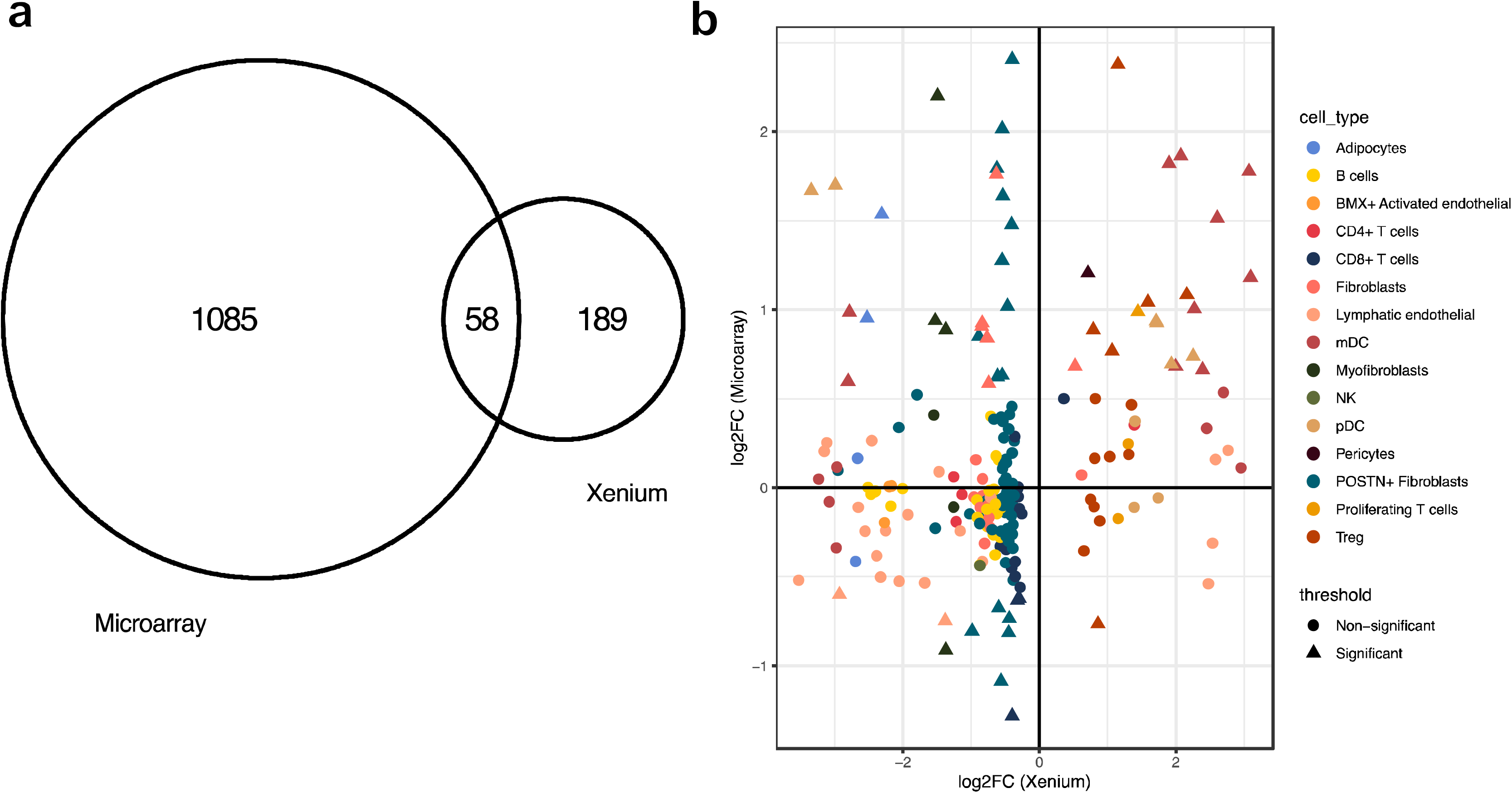
**(A)** Venn diagram showing overlap between DEGs (adjusted P-value < 0.05) in Xenium compared to DEGs (Benjamini-Hochberg corrected P-value < 0.05 and |log_2_FC| > 1.5) in a publicly-available EMB microarray data set (GSE150059) comparing 76 patients with ACR and 179 patients with AMR. **(B)** Scatter plot of the 247 unique genes that were DE between ACR and AMR in our study (Xenium). The x-axis represents log_2_FC in the Xenium data set while the y-axis represents log_2_FC in the microarray data set. Significant genes, depicted by triangles, are significant in both data sets as described above. Colors represent cell type specificity of DE genes from the Xenium data set. When more than one cell type-gene pair was significant in the Xenium data set, the cell type with the lowest adjusted P-value is depicted.

**Extended data Fig. 3.**
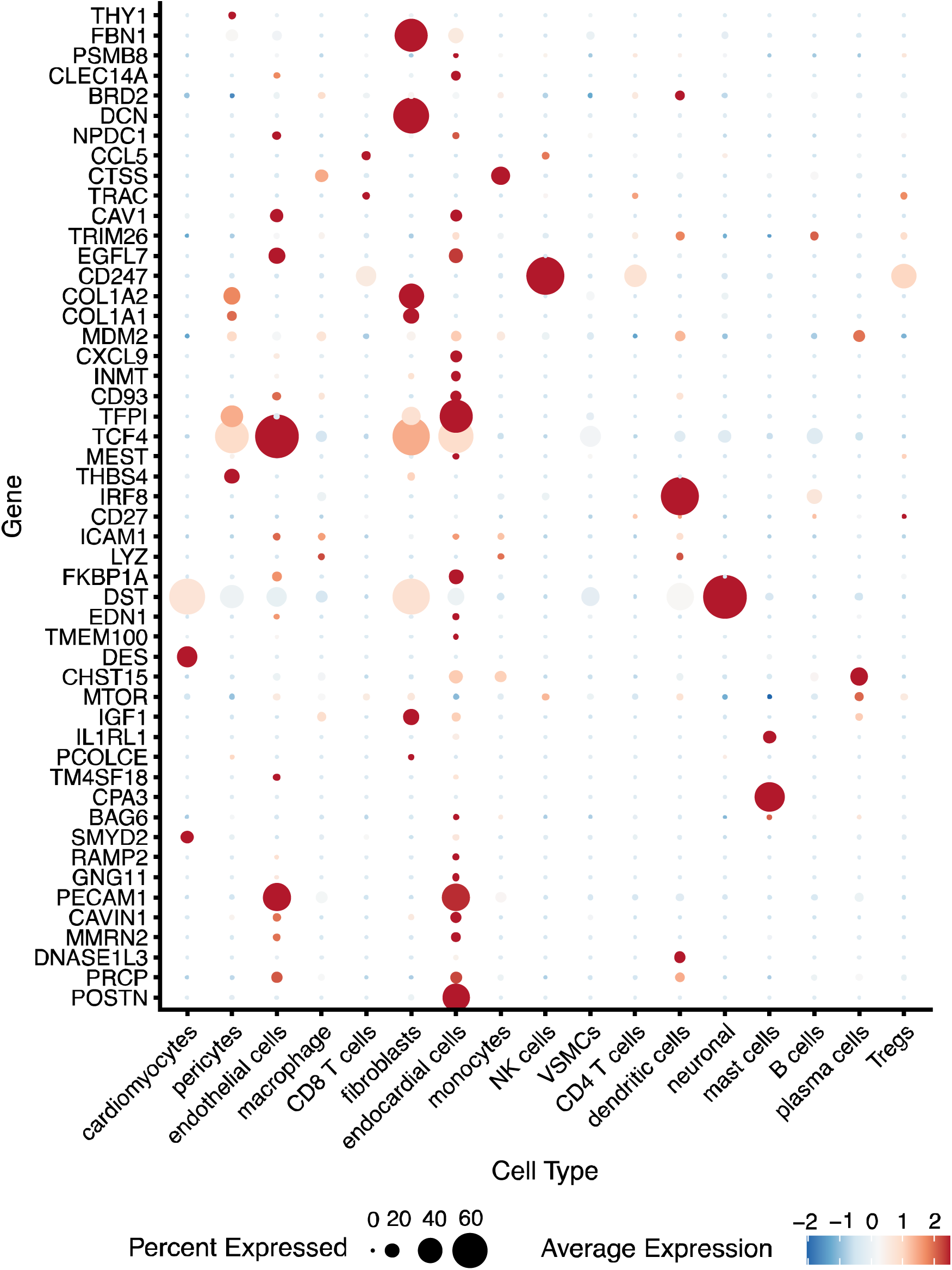
Dot plot of the same top 50 unique genes from **Fig 5C** showing expression in a single-nuclear RNA-sequencing data set consisting of right ventricular cardiac biopsies from patients with end-stage CAV requiring re-do heart transplantation.

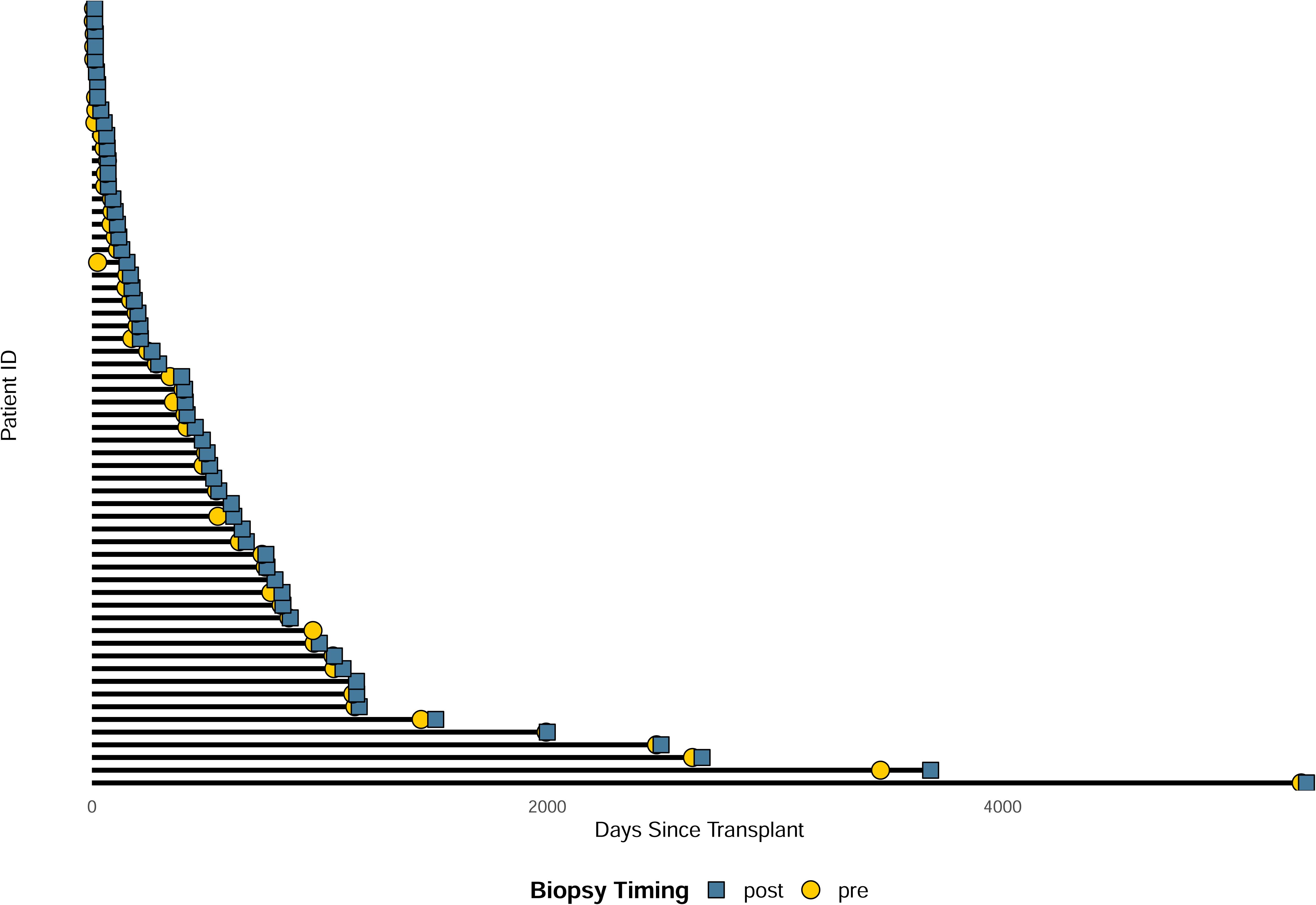

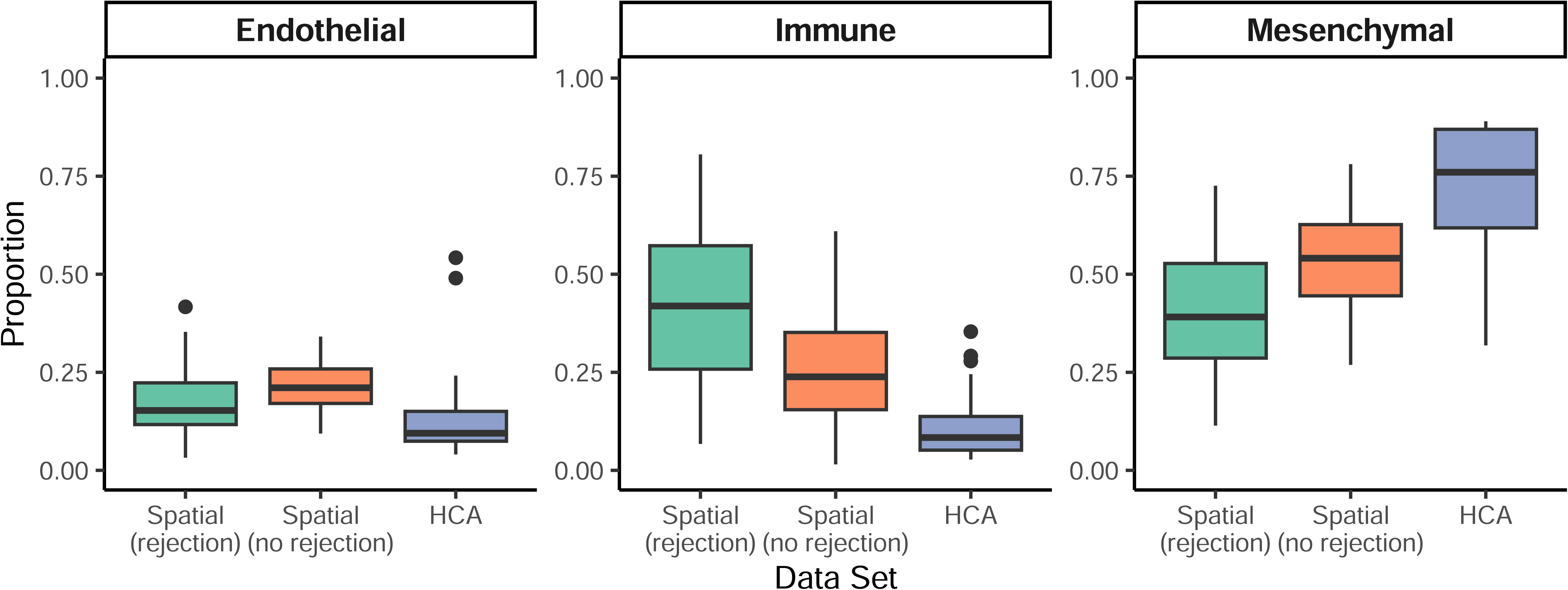

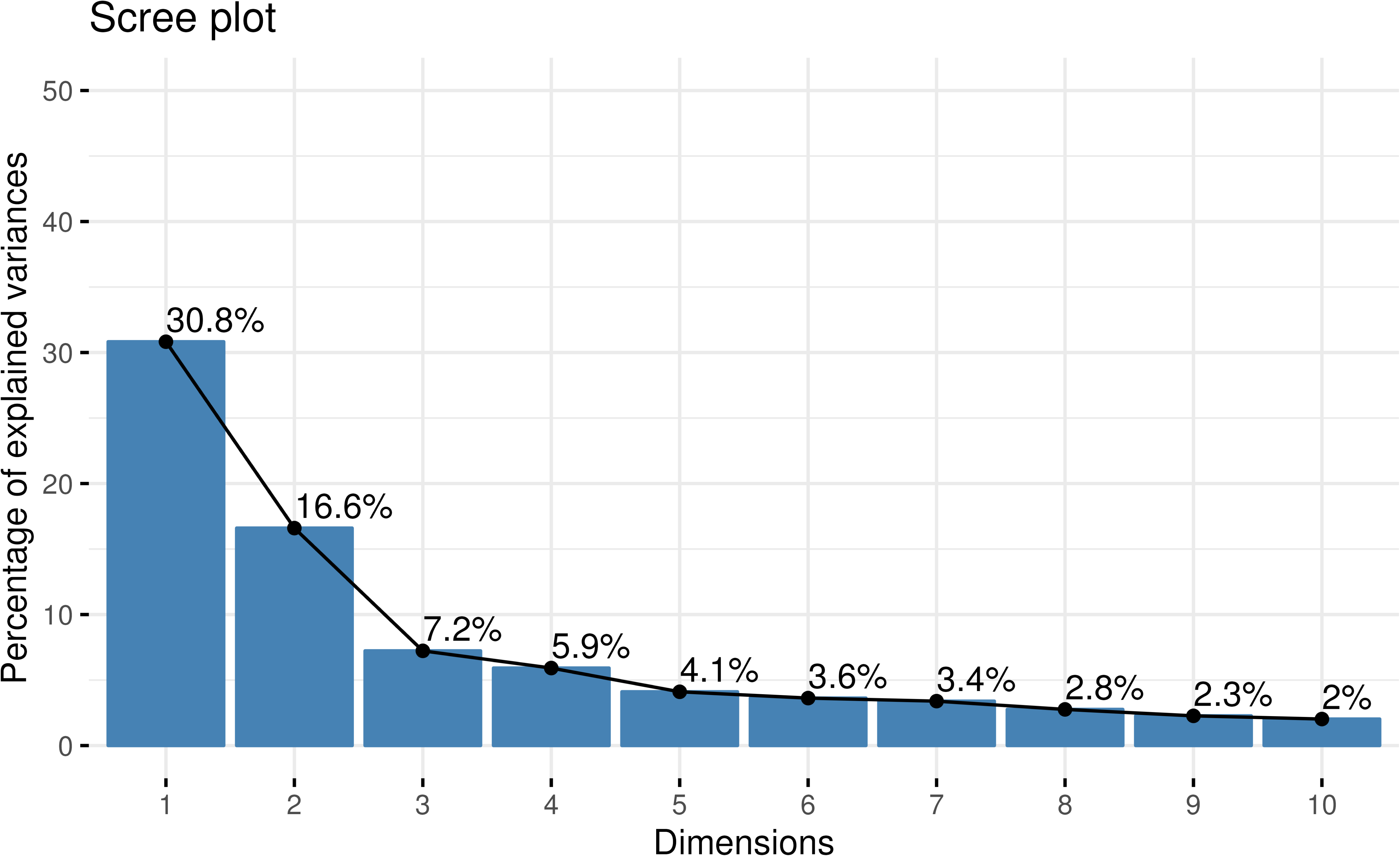

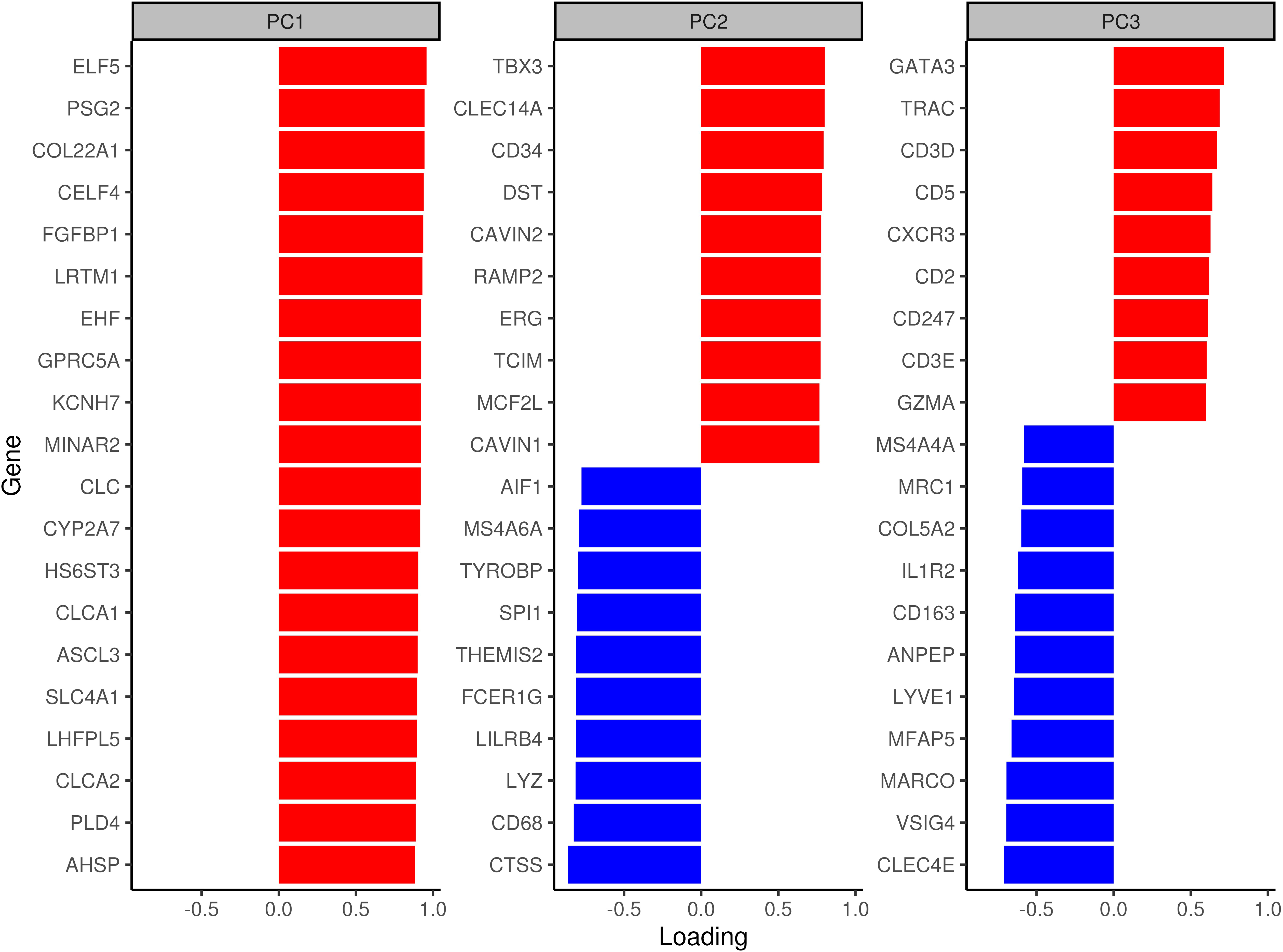

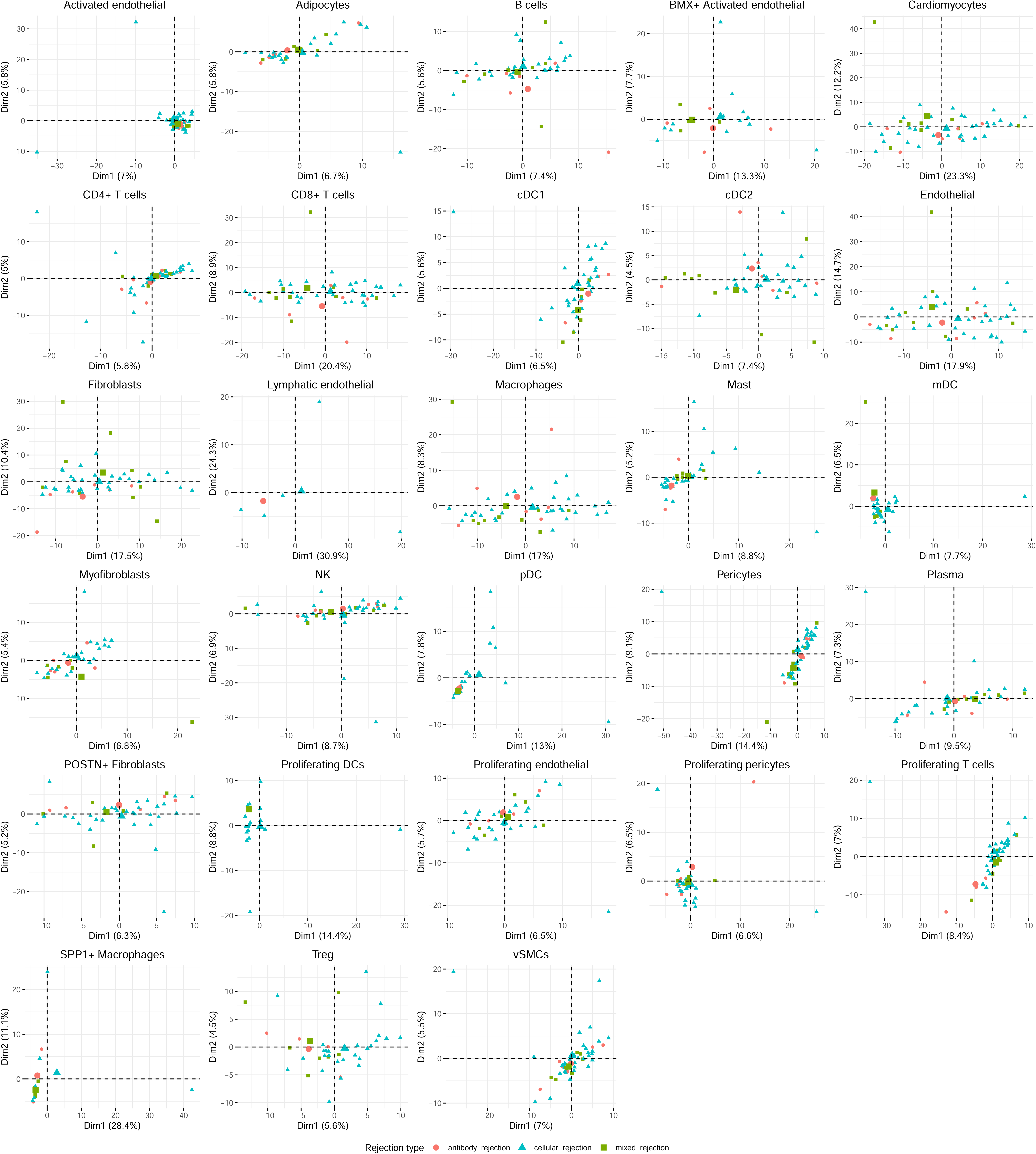

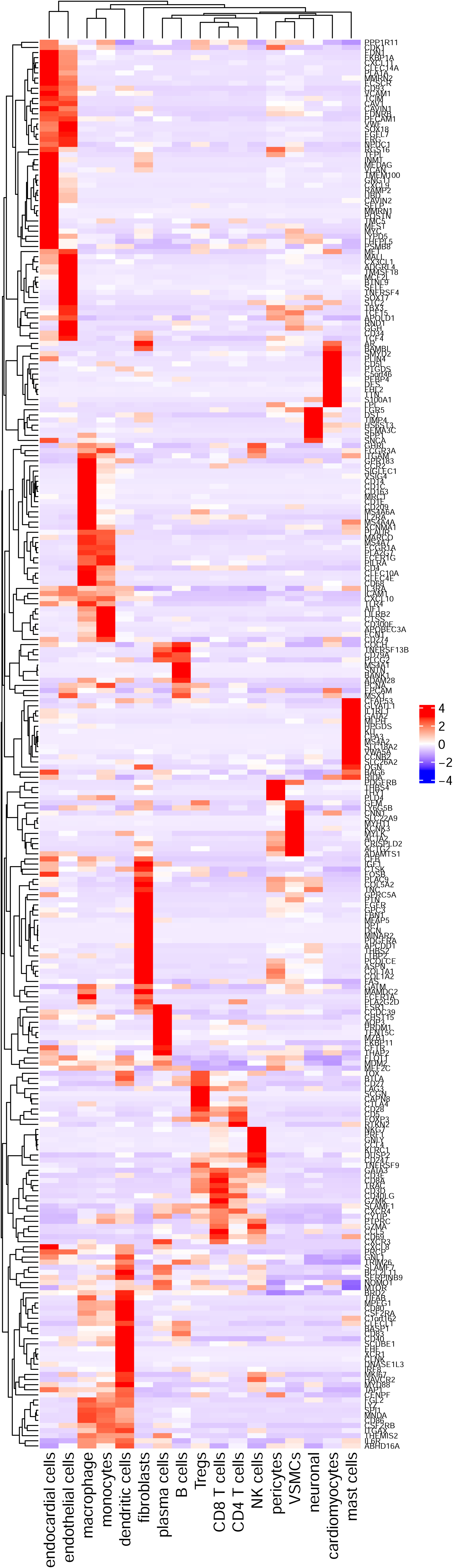

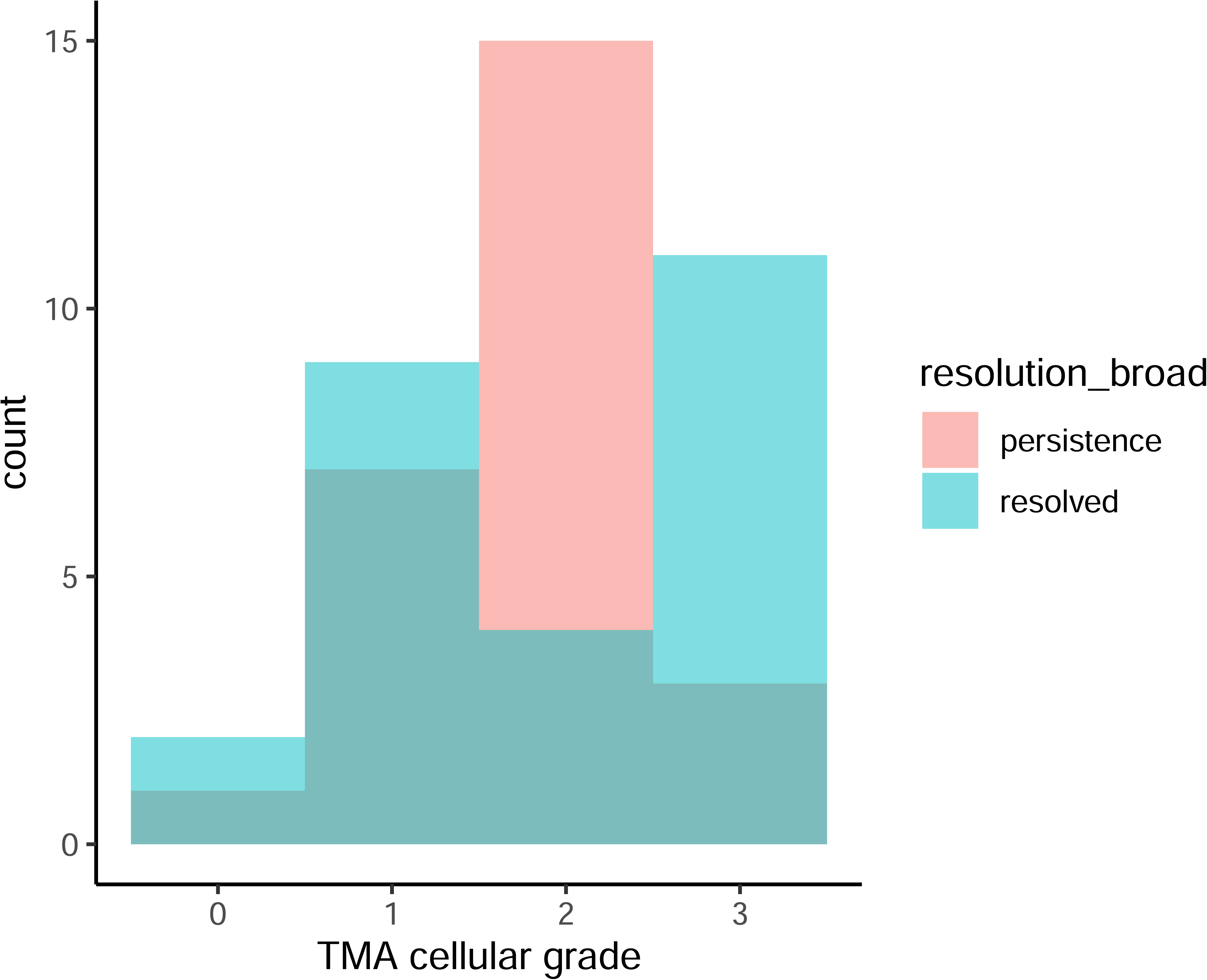

