## Supplemental Material for "Dynamic responses to rejection in the transplanted human heart revealed through spatial transcriptomics"

**SUPPLEMENTARY FIGURES**

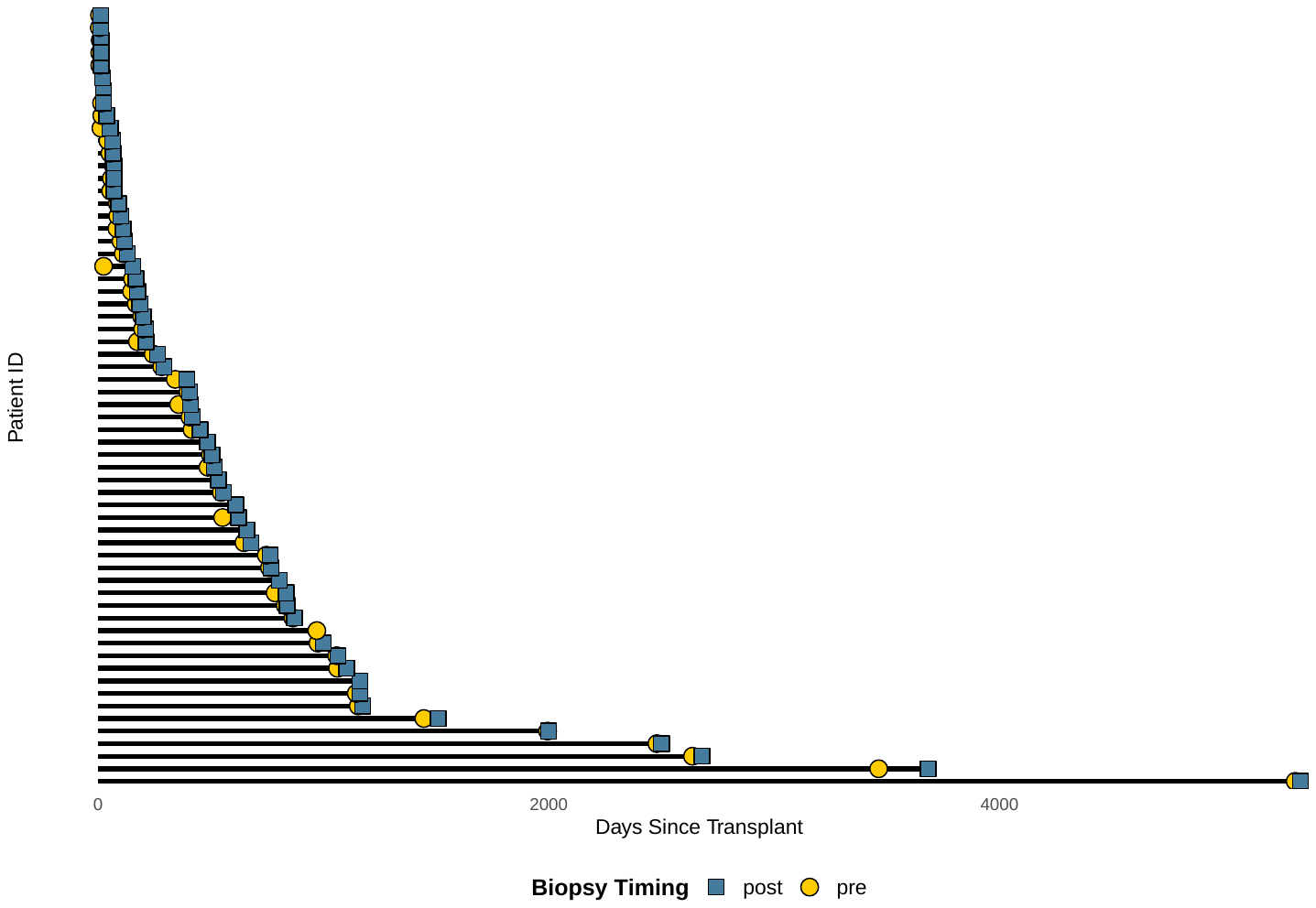
**Supplementary Figure 1.** Swimmer’s plot of time from transplant to biopsy (in days).

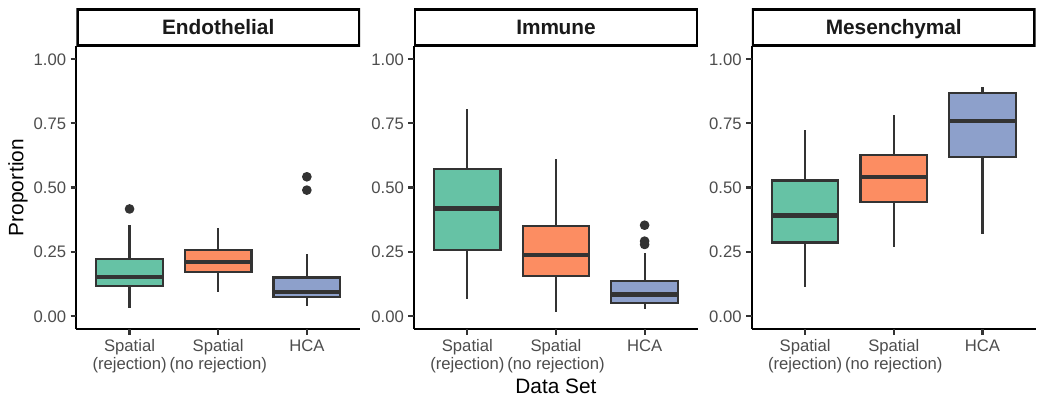
**Supplementary Figure 2.** Comparison of cell-type recovery between the present spatial dataset and scRNA data set from the Human Heat Cell Atlas (HCA). Proportion of cells for each lineage were calculated for rejection biopsies and non-rejection biopsies in the present spatial data set while the proportion of cells from each lineage was calculated for each donor in the HCA data.

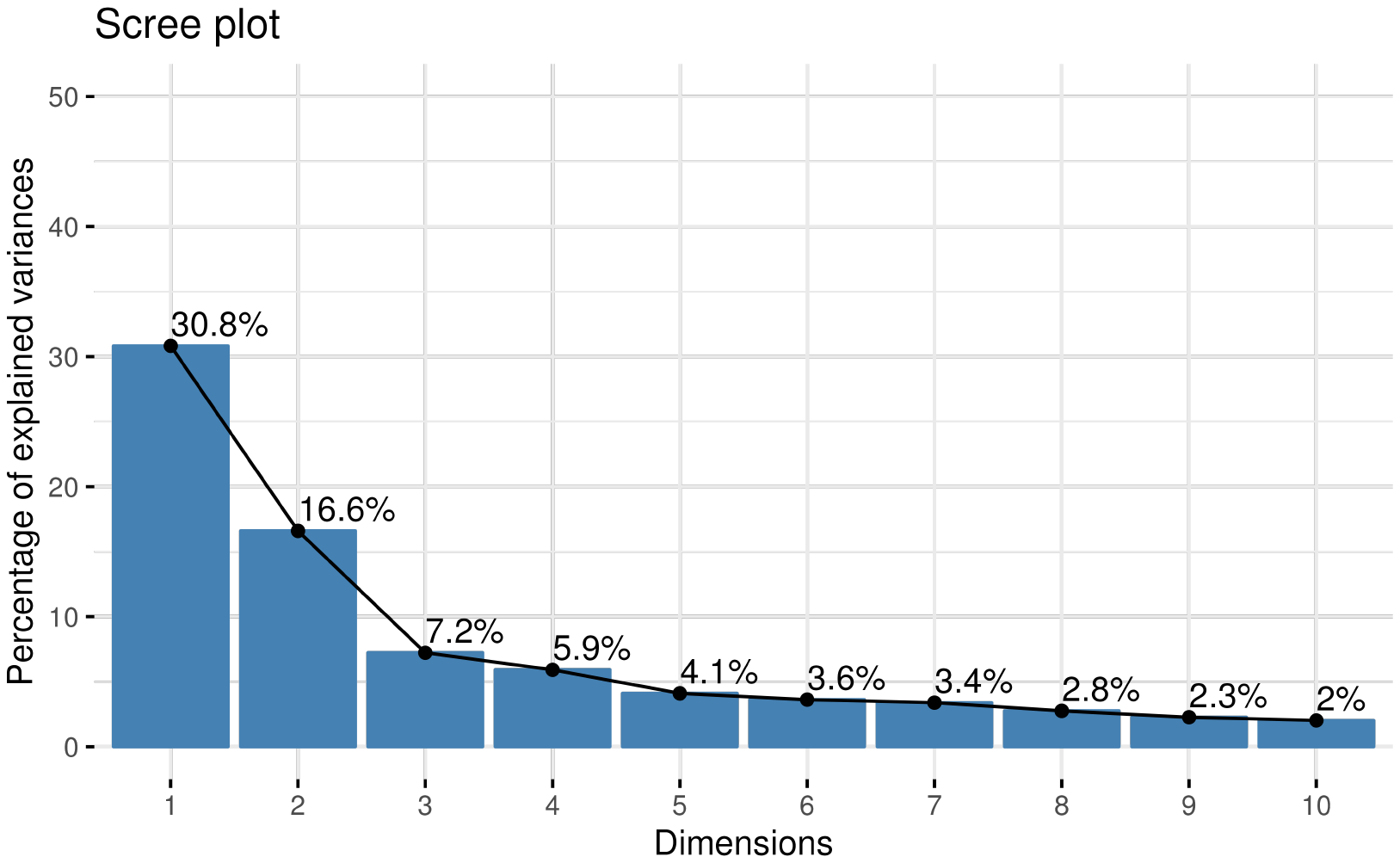
**Supplementary Figure 3.** Scree plot shows that 54.6% of variation is contained within the first 3 principal components (PCs), which were carried forward for downstream analyses.

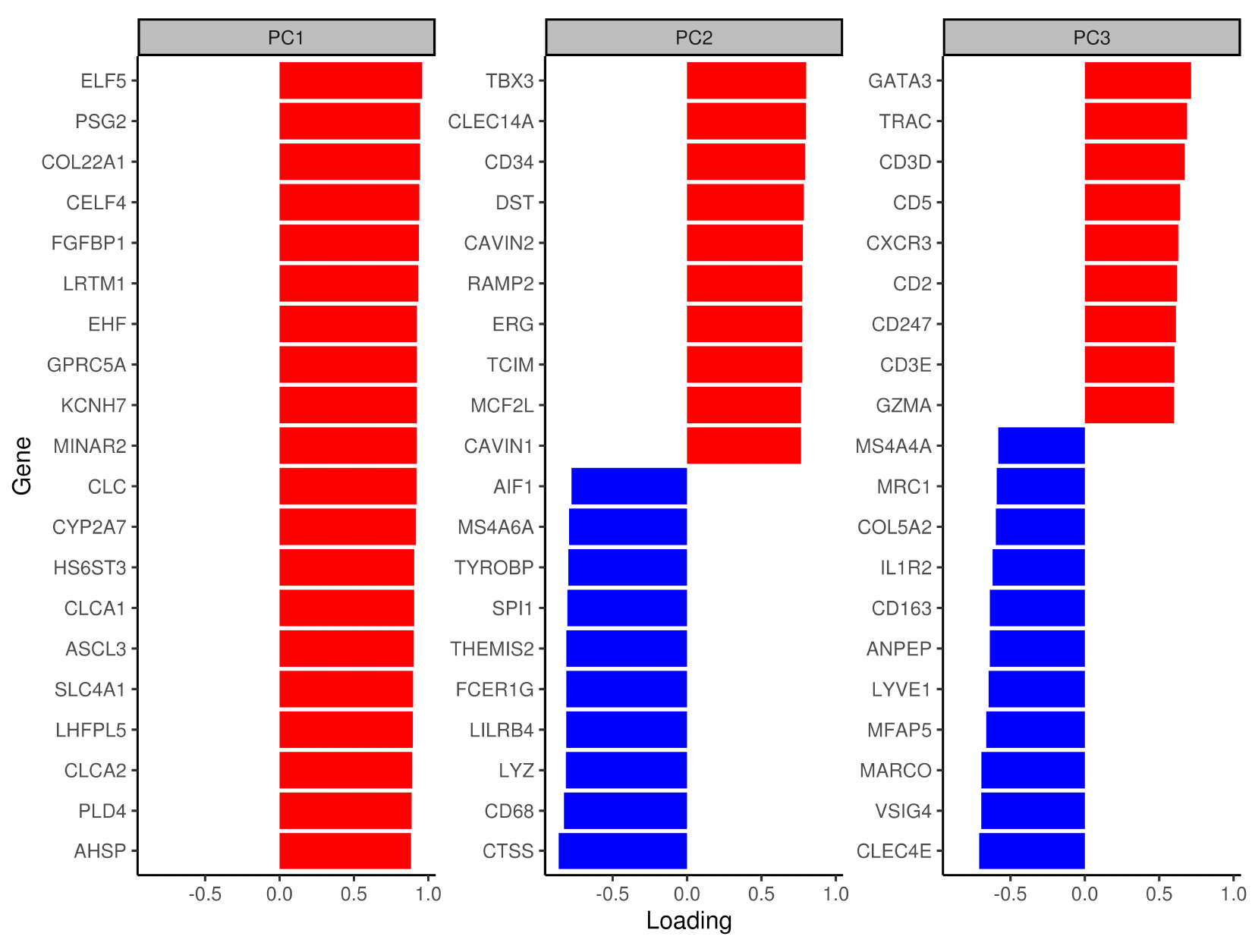
**Supplementary Figure 4.** PC loadings of the top 20 genes by their absolute loadings in each of the first 3 PCs.

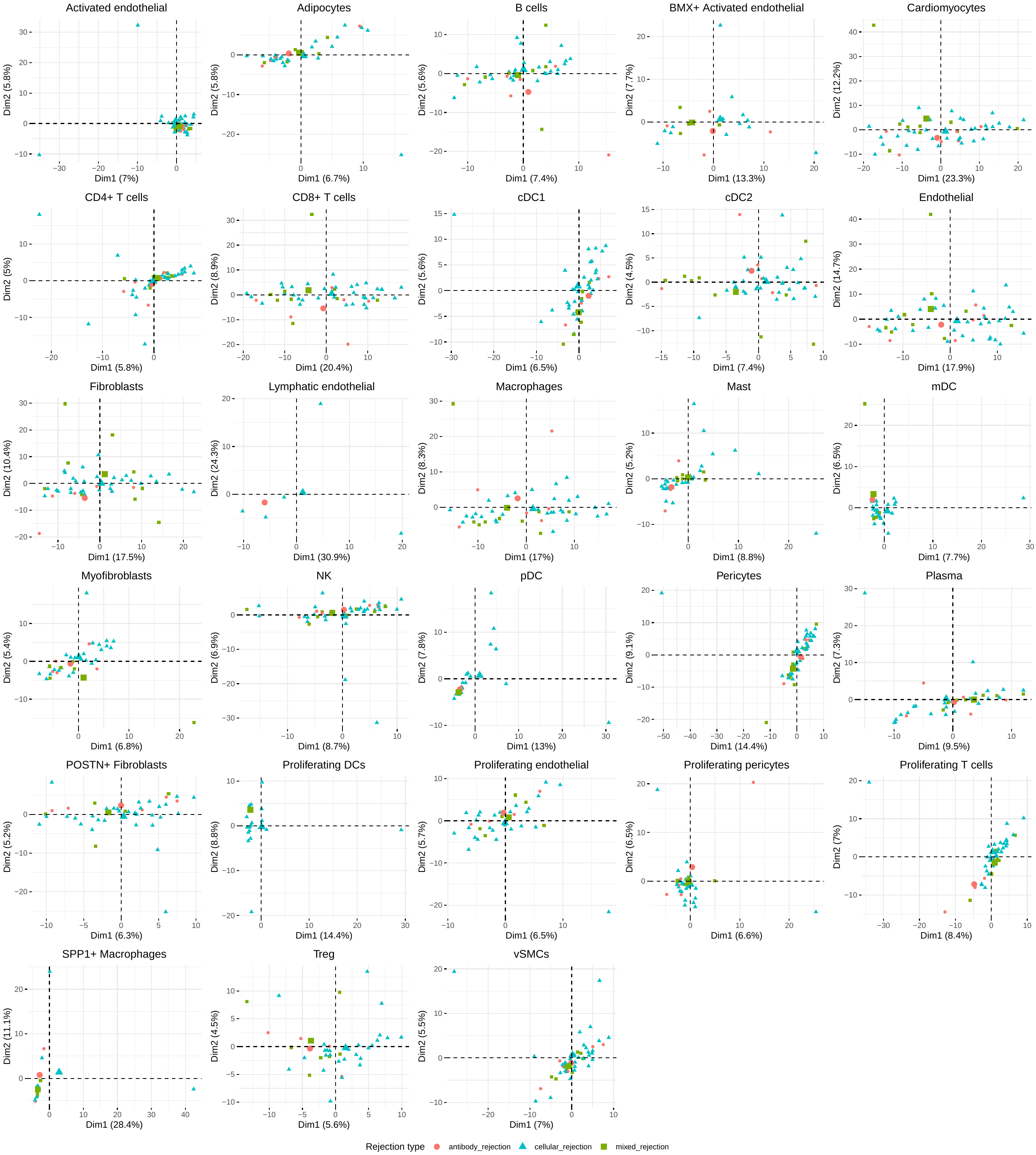
**Supplementary Figure 5.** Cell type-specific PCA plot for all 28 cell types based on gene expression patterns spanning all three rejection types.

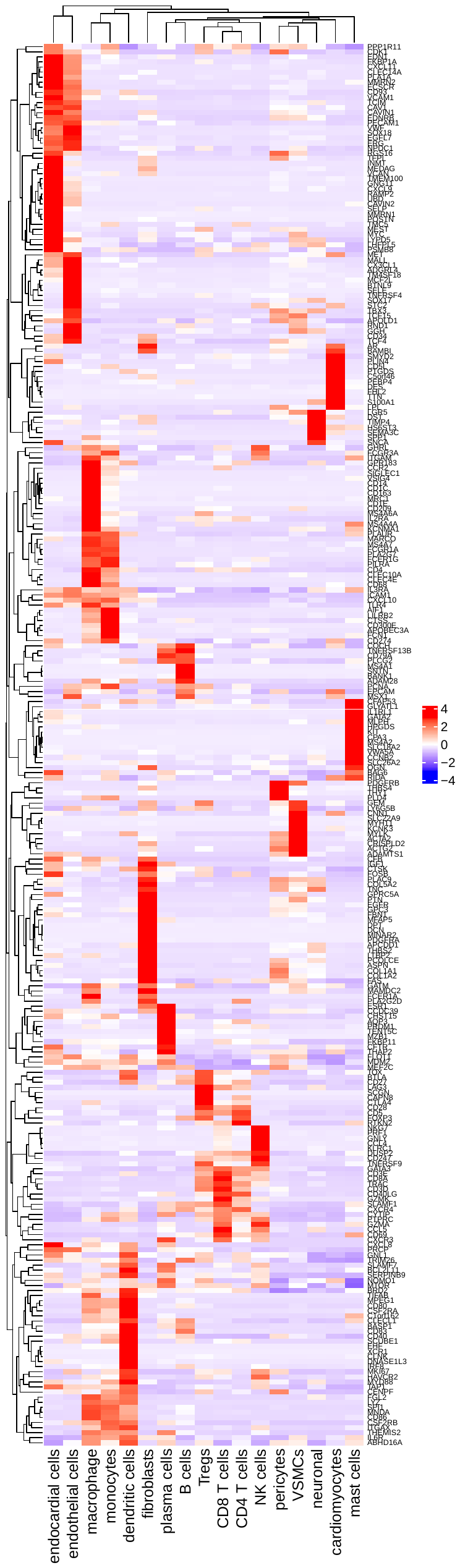

**Supplementary Figure 6.** Heatmap of all 309 unique genes that were significantly associated with CAV in our study (Xenium) depicted in an external single-nuclear RNA-sequencing data set of cardiac biopsies from the right ventricle in patients with end-stage CAV requiring re-do heart transplantation. Normalized expression by cell type is shown.

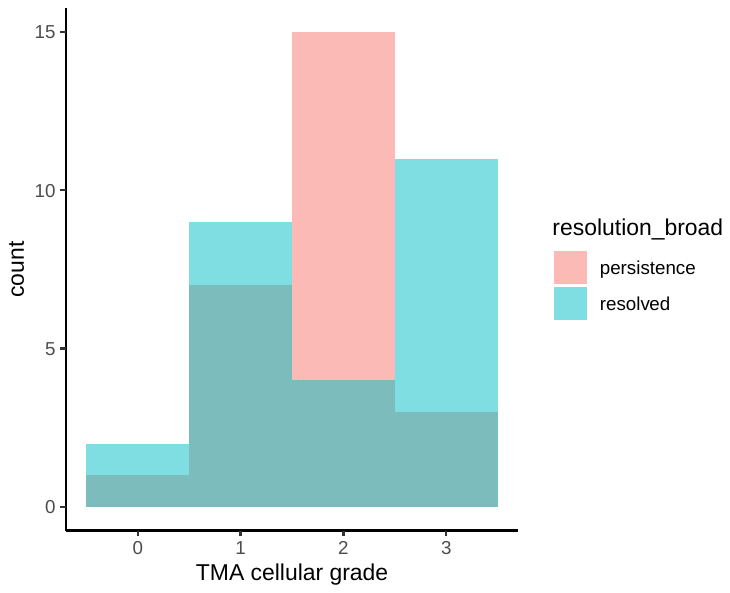

**Supplementary Figure 7.** Distribution of TMA grade for ACR grade 2R biopsies between patients whose rejection resolved (responders) and those whose rejection persisted (nonresponders) despite immunomodulatory therapies.
